# FEZ1 participates in human embryonic brain development by modulating neuronal progenitor subpopulation specification and migration

**DOI:** 10.1101/2022.07.11.499073

**Authors:** Yinghua Qu, Omer An, Henry Yang, Yi-Chin Toh, John Chua Jia En

**Affiliations:** Department of Physiology, Yong Loo Lin School of Medicine, National University of Singapore, Singapore; Department of Biomedical Engineering, National University of Singapore, Singapore; Cancer Science Institute of Singapore, National University of Singapore, Singapore; School of Mechanical, Medical and Process Engineering, Queensland University of Technology, Australia; Centre for Biomedical Technologies, Queensland University of Technology, Australia; Healthy Longevity Translational Research Program, Yong Loo Lin School of Medicine, National University of Singapore; Neurobiology/Ageing Programme, National University of Singapore, Singapore; Institute for Molecular and Cell Biology, A*STAR, Singapore

## Abstract

Abnormal neuronal networks arising from perturbations during early brain development contribute to neurodevelopmental disorders. Mutations and deletions of human Fasciculation and Elongation Protein Zeta 1 (*FEZ1*) are found in schizophrenia and Jacobsen syndrome patients. However, its roles in human brain development and manifestation of clinical pathological symptoms remain unknown. Here, using human cerebral organoids (hCOs), we observed that *FEZ1* expression is turned on early during brain development and is detectable in both neuroprogenitor subtypes and immature neurons. Deletion of *FEZ1* disrupts expression of genes involved in neuronal and synaptic development. Using single-cell RNA sequencing, we further uncovered an abnormal expansion of homeodomain-only protein homeobox (HOPX)^−^ outer radial glia (oRG) in FEZ1-null hCOs, occurring at the expense of HOPX^+^ oRG. HOPX^−^ oRGs show higher cell mobility as compared to HOPX^+^ oRGs, which is accompanied by the ectopic localization of the neuroprogenitors to the outer layer of FEZ1-null hCOs. Moreover, abnormal encroachment of TBR2^+^ intermediate progenitors into CTIP2^+^ deep layer neurons indicated that cortical layer formation is disrupted in FEZ1-null hCOs. Collectively, our findings highlight the involvement of FEZ1 in early cortical brain development and how it contributes to neurodevelopmental disorders.

## INTRODUCTION

Cortical development is a multistage process involving neurogenesis, cell migration, differentiation and maturation that occurs over an extended time frame (Klingler et al., 2021; Lui et al., 2011; Paridaen and Huttner, 2014; Silbereis et al., 2016). Despite a wealth of knowledge in the identities of transcription factors driving neurogenesis and differentiation of neural progenitors to different neuronal subtypes, the compendium of other genes participating in the various developmental stages is only beginning to be progressively unravelled. Significantly, mutations in a number of them have been associated with human brain disorders with neurodevelopmental origins, including neuropsychiatric disorders such as schizophrenia (SCZ), attention deficit hyperactivity disorder (ADHD) and autism spectrum disorders (ASD) (Gandal et al., 2018; Rosato et al., 2019; Walker et al., 2019). Nevertheless, how these genes contribute to such disorders remain incompletely understood.

One such implicated gene is Fasciculation elongation protein zeta 1 (FEZ1). Its invertebrate homolog, *Unc-76,* was initially identified as necessary for nervous system development in *Caenorhabditis elegans. unc-76* mutants showed axonal outgrowth and fasciculation defects as well as abnormalities in synapse formation and organization (Bloom and Horvitz, 1997; Butkevich et al., 2016; Desai et al., 1988; Su et al., 2006). These phenotypes were also recapitulated in *Drosophila unc-76* mutants (Gindhart et al., 2003; Toda et al., 2008). Importantly, deletion of FEZ1 in mammalian neurons severely retarded outgrowth and branching of axons and dendrites, suggesting that impaired neurodevelopment as a result of FEZ1 loss can contribute to neurodevelopment and neuropsychiatric disorders (Chua et al., 2021; Gunaseelan et al., 2021; Kang et al., 2011). Supporting this notion, FEZ1-knockout mice exhibited behavioural defects akin to those observed in SCZ and ADHD patients (Sakae et al., 2008; Sumitomo et al., 2018).

Significantly, *FEZ1* polymorphisms and changes in brain expression of *FEZ1* have been detected in SCZ patients (Colantuoni et al., 2008; Kang et al., 2011; Lipska et al., 2006; Tang et al., 2017; Vachev et al., 2015; Yamada et al., 2004). The gene is also frequently lost in Jacobsen syndrome patients, a rare disorder affecting children where the terminal region of chromosome 11q is deleted (Favier et al., 2015; Grossfeld et al., 2004; Mattina et al., 2009). These patients are frequently diagnosed with ADHD, while ASD, SCZ and rarely, bipolar disorders have also been documented. These observations point towards *FEZ1*’s importance in human brain development and its dysfunction as contributing to these disorders. However, its roles in human cortical brain development and its involvement in neurodevelopmental disorders have not been formally established.

Here, using human cerebral organoids (hCO) as a human brain development model, we observed that *FEZ1* mRNA and protein, which was absent in human embryonic stem cells (hESCs), became detectable in neuroprogenitor cell types early during hCO development and continued to increase as the organoids matured. While loss of FEZ1, engineered by CRISPR-Cas9 gene-editing, did not cause significant morphological changes in FEZ1-null hCOs, RNAseq analyses uncovered a substantial number of differentially expressed genes enriched in neuronal development as well as synaptic function, supporting previous studies demonstrating its importance in these biological processes. scRNA-seq analyses of hCOs further uncovered an abnormal expansion of homeodomain-only protein homeobox negative (HOPX^−^) outer radial glial (oRG) in FEZ1-null hCOs that came at the expense of a diminished pool of HOPX^+^ canonical oRG. Analyses of differentially expressed genes (DEGs) between HOPX^−^ oRG and HOPX^+^ oRG indicated differences in expression of genes involved in actin dynamics. Supporting this, oRGs in FEZ1-null hCOs showed higher globular (G-) to filamentous (F-) actin ratios and anomalous cell migration *in situ.* These changes were accompanied by a corresponding increase and anomalous expansion of TBR2^+^ intermediate progenitors (IP) into a reduced CTIP2^+^ layer of deep layer neurons in FEZ1-null hCOs, resulting in early cortical lamination abnormalities. Taken together, our results supported earlier findings of FEZ1’s roles in neuronal development and synapse formation and additionally uncovered an unexpected involvement of FEZ1 in influencing developmental trajectories and cortical layer formation during cortical brain development.

## Results

### Expression of FEZ1 begins early during human brain development

Rare mutations and deletions in *FEZ1* have been linked to human neurodevelopmental disorders such as schizophrenia and Jacobsen syndrome (Jurcă et al., 2017; Kang et al., 2011; Vachev et al., 2015; Yamada et al., 2004). However, its role in human brain development is unknown. In rodents, detection of *FEZ1* mRNA and protein coincided with the height of neurogenesis, indicating its likely involvement in the earliest phases of brain development (Fujita et al., 2004; Sakae et al., 2008). To examine whether this might occur during human brain development, we generated human cerebral organoids (hCOs) following a non-patterning factor based protocol (Lancaster and Knoblich, 2014). As reported, rosette-like structures with apical ventricular zones (VZ) surrounded by PAX6^+^ neuroepithelial cells appeared in developing hCOs as early as day 10 (D10) (Fig. S1, A and B) (Lancaster and Knoblich, 2014; Qian et al., 2016). At D28, sequential layering of TBR2^+^ intermediate progenitors (IP) and CTIP2^+^ deep layer neurons can be observed, indicating that cortical layering has begun at this time, which concurred with earlier reports (Fig. S1B) (Lancaster and Knoblich, 2014; Qian et al., 2016).

Consistent with the early expression of FEZ1 in rodent brain development, FEZ1 expression was apparent as cytosolic puncta within PAX6^+^ neuroepithelial cells in the at VZ regions of D10 hCOs (Fig. 1A). By D28, both FEZ1 levels and the proportion of PAX6^+^ cells expressing FEZ1 had further increased. Moreover, its expression expanded from VZ to include the subventricular (SVZ) and preplate (PP) (Matsumoto et al., 2020; Pollen et al., 2019)(Fig. 1A). By D60, FEZ1 puncta could be observed in TUJ1^+^ neurites emanating from newly derived neurons in the cortical plate (CP) (Mora-Bermudez et al., 2016; Qian et al., 2016)(Fig. 1A). Immunoblot and real-time PCR analyses further confirmed that FEZ1 expression, while virtually undetectable in undifferentiated human embryonic stem cells (hESCs), progressively increased as hCOs developed (Fig. 1, B and C). To further ascertain that FEZ1 expression was switched on at the earliest stages of cortical development, we directly derived human neuroepithelial cells (hNEs) from hESCs (Ying et al., 2003). As observed in hCOs, increased levels of both *FEZ1* mRNA and protein followed differentiation of hESCs (Fig. 1, D and E) and human induced pluripotent stem cells (hiPSCs) (Fig. S1C) to NEs. Taken together, our results indicated that FEZ1 expression began at the early stages of neural specification independently of the induction protocol or culture configuration, supporting its involvement in early human brain development.

**Figure 1.**
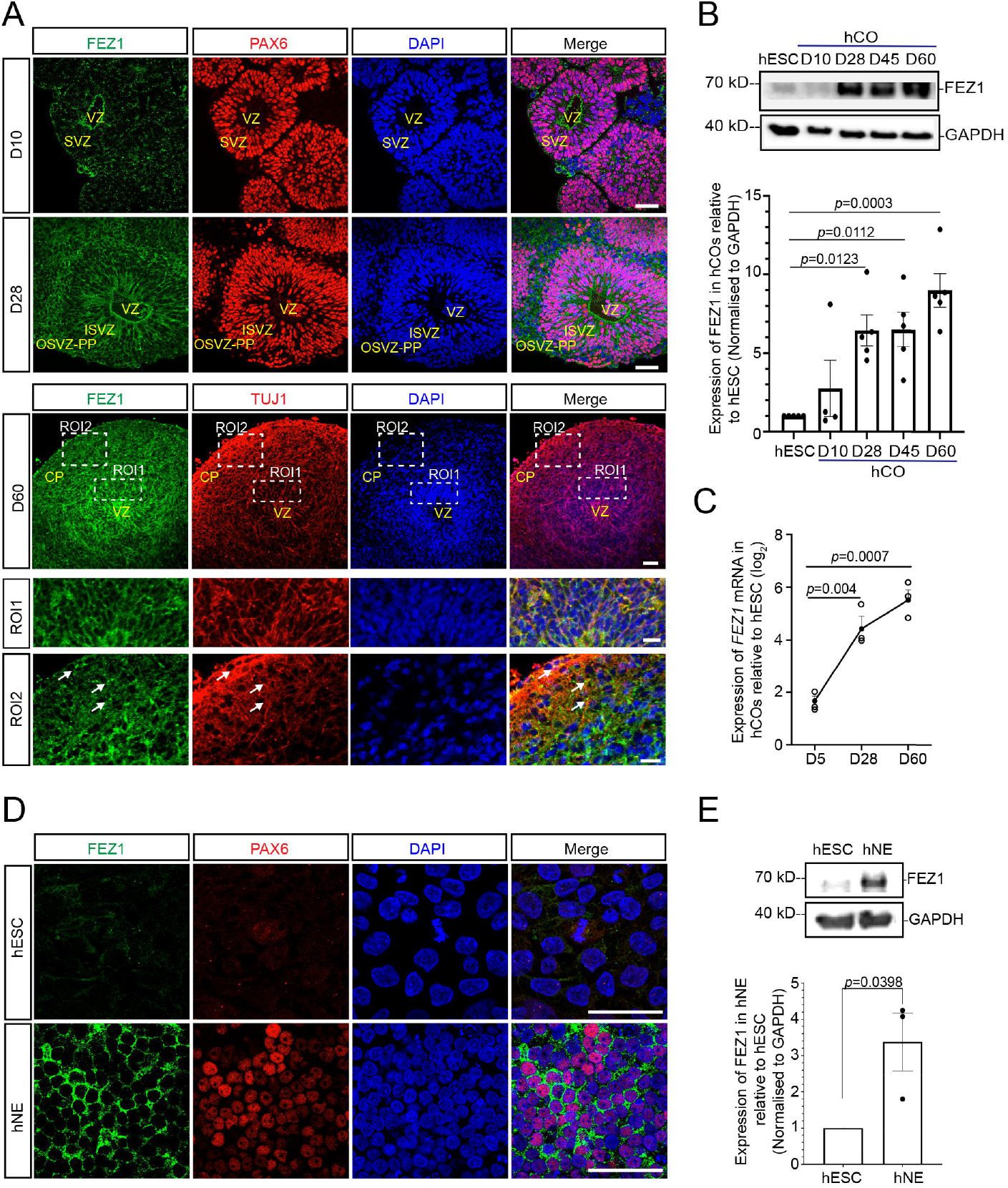
FEZ1 expression increased progressively during human cerebral organoid (hCO) development. (A) Immunofluorescence (IF) staining of FEZ1 with the hNE marker PAX6 in D10, D28 and D60 hCOs (upper panel). Co-staining of FEZ1 with the neuronal marker TUJ1 in D60 hCOs (lower panel). Region of interests (ROIs) show higher magnification of the ventricular zone (VZ), sub-ventricular zone (SVZ) and cortical plate (CP) in D60 hCOs. Punctate FEZ1 staining is visible. In ROI2, arrows indicate FEZ1 puncta along TUJ^+^ neurites in the OSVZ. Scale bars, 50 μm (ROI1) and 20 μm (ROI2). (B) Immunoblot and quantification of FEZ1 protein expression during hCO development. Values represent mean ± SEM (from n=5 independent organoid differentiations, ordinary one-way ANOVA with Tukey’s multiple comparisons test with p values indicated). (C) RT-qPCR analysis of FEZ1 mRNA expression in D5 to D60 hCOs. Results were normalized against GAPDH and expressed relative to FEZ1 levels in undifferentiated human embryonic stem cells (hESCs). Values represent mean ± SEM (n=3 independent organoid differentiations, ordinary one-way ANOVA with Tukey’s multiple comparisons test with p values indicated). (D) IF staining of FEZ1 in hESC and hNE shows that FEZ1 expression is switched on upon differentiation of hESC to human neuroepithelium (hNE). Scale bars, 50 μm. (E) Immunoblot and quantification of FEZ1 protein expression in hESC (H1) and hNE. Values represent mean ± SEM (n= 3 independent hESC to hNE differentiations, unpaired t-test determined the two-tailed *p* value as indicated). VZ: ventricular zone; SVZ: sub-ventricular zone; ISVZ: inner sub-ventricular zone; OSVZ-PP: outer sub-ventricular zone-preplate.

### Loss of FEZ1 alters expression of genes involved in early brain and neuronal development

To examine the role of FEZ1 in early human brain development, we generated FEZ1-null hESCs using a lentivirus-mediated CRISPR-Cas9 strategy that targeted exon 2 of the *FEZ1* gene (Fig. 2A) (Chua et al., 2021; Gunaseelan et al., 2021). Following puromycin selection, 24 clones were independently selected for expansion. Of these, 3 clones were further examined by amplification and sequencing of the targeted genomic region to confirm the presence of indels in the targeted region. All 3 clones contained an identical insertion of an adenine residue at position 366 in the mRNA sequence. This introduced a frameshift in the open reading frame of the coding region, giving rise to a prematurely truncated peptide containing the first 58 amino acids of FEZ1, followed by the addition of 6 non-templated amino acids (Fig. 2A).

**Figure 2.**
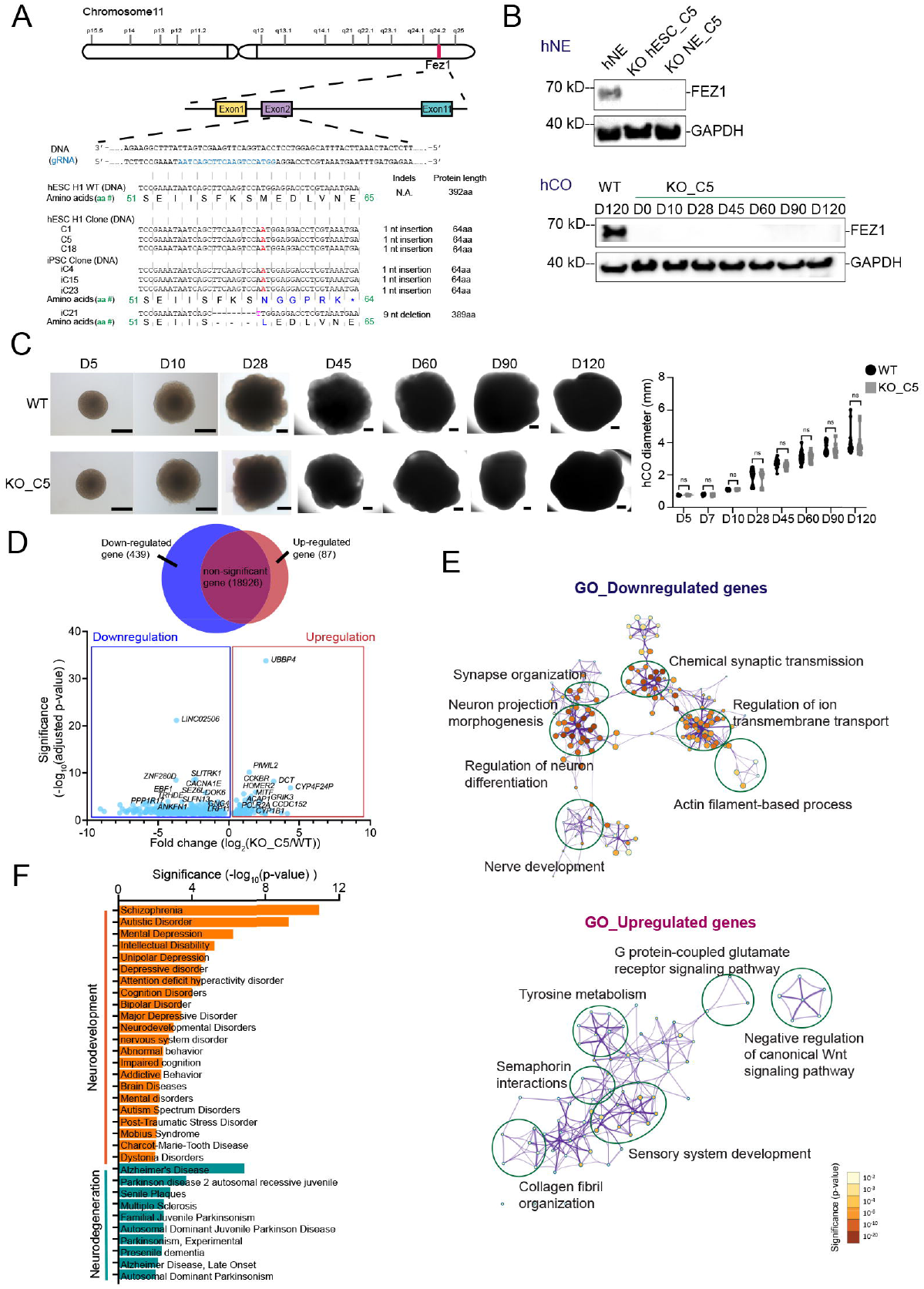
Deletion of FEZ1 altered expression of genes involved in neurodevelopment and neurodevelopmental disorders. (A) CRISPR-Cas9-mediated FEZ1 deletion in hESCs and hiPSCs. All clones selected from both hESC (H1) and hiPSC lines consistently harbored a 1 bp insertion (shown in red) in exon 2. Only one hiPSC clone (iC21) contained a 9 bp deletion (shown in magenta). (B) Immunoblot of FEZ1 expression in hNE and developing hCOs confirmed the successful ablation of FEZ1 expression in FEZ1-null hESC_C5 clone. (C) hCOs generated from wildtype (WT) hESC H1 line and FEZ1-null hESC_C5 clone showed no gross differences in morphology and size over 120-day culture period. Scale bar: 500 μm. (n=4 independent organoid differentiations for WT and FEZ1-null group, respectively. Each data point represents measured diameter of one organoid, at least 4 organoids were measured in each replicate, two-way ANOVA with Šidák’s multiple comparisons, ns: not statistically significant). (D) Venn diagram (top panel) and volcano plot (bottom panel) of significant differentially expressed genes (DEGs) (log_2_FC > 1 or < −1; adjusted p-value < 0.05) between D10 FEZ1-null hCO and D10 WT hCOs. Top 25 DEGs are shown in the plot. (E) Gene Ontology (GO) analysis of significant DEGs in D10 FEZ1-null hCOs using Metascape (https://metascape.org/gp/index.html#/main/step1). Statistical significances of enriched GO terms are shown in the color-coded legend. (F) Disease enrichment analysis of significant DEGs in D10 FEZ1-null hCOs using Enricher (https://maayanlab.cloud/Enrichr/). DEGs in FEZ1-null hCOs were significantly associated with neurodevelopment and neurodegeneration disorders.

In parallel, we also generated independent FEZ1-null hiPSC lines using the same strategy. Four independent clones were obtained and analyzed by DNA sequencing as before. Three of these clones contained identical frameshift insertions present in the hESC clones (Fig. 2A). The last clone (iC21) contained 10 nucleotide deletion followed by a single base pair in-frame insertion within the coding region (position 356-375). This resulted in the predicted translation of a full-length FEZ1 without its 3 amino acids at positions 56-58 and the substitution of methionine at position 59 with leucine. Since most of the N and C terminal functional motifs were preserved, iC21 was not further studied.

Immunoblot analyses of cell lysates from monolayer NEs and hCOs derived from 2 selected FEZ1 null clones (KO hESC_C5 and KO hiPSC_iC4) confirmed successful elimination of FEZ1 expression (Fig. 2B; Fig. S2A). KO hESC_C5 (henceforth called FEZ1-null hESC) was used as the representative FEZ1-null H1 hESC clone in all subsequent studies since all FEZ1-null hESC and iPSC clones possessed identical frameshift insertion site in *FEZ1* gene. We further determined the absence of gross chromosomal abnormalities in this clone after gene editing by karyotyping (Fig. S2B). Additionally, we did not detect significant differences in cell proliferation rate or the expression of the pluripotency markers SOX2, LIN28, NANOG and SSEA4 (Fig. S2C-D). This indicated that loss of *FEZ1* did not cause adverse effects on stem cell pluripotency or survival.

Macroscopically, loss of FEZ1 did not appear to affect generation of hCOs from the FEZ1-null hESC. There were no significant differences in gross morphology or size of hCOs derived from wildtype and FEZ-null hESCs over a culture period of 120 days (Fig. 2C). Moreover, immunofluorescent staining of both groups of hCOs at D10 with Pax6 and Nestin shows the formation of rosettes by neuroepithelial cells (Fig. S2E). Supporting this, no difference in size was also observed between hCOs derived from wildtype and FEZ1-null hiPSC up to 28 days (Fig. S2F). However, FEZ1-null hCOs derived from KO-iC4 iPSCs became somewhat smaller from D45 onwards.

Although deletion of FEZ1 gene did not induce gross aberrations in hCO formation and morphology, it could still cause changes to cellular signaling and programming pathways. Indeed, bulk RNA-seq analyses of wildtype and FEZ1-null hCOs at D10 uncovered significant changes between the 2 transcriptomes. In total, 526 differentially expressed genes (DEGs) were identified (Fig. 2D; Table S1). Of these, 87 genes were up-regulated and 439 genes were down-regulated in FEZ1-null as compared to wildtype hCOs (Fig. 2D). Gene ontology (GO) enrichment analysis revealed that genes involved in neuronal development and synaptic function were significantly over-represented in the down-regulated genes (Fig. 2E, upper panel; Table S2). These results correlated well with the established function of FEZ1 in neuronal development and synaptic function (Butkevich et al., 2016; Chua et al., 2021; Dong et al., 2021; Gunaseelan et al., 2021; Kang et al., 2011). Genes upregulated in the absence of FEZ1 were primarily associated with Wnt signaling, sensory system development, tyrosine metabolism and semaphorin interactions (Fig. 2E, bottom panel; Table S3). Wnt and semaphorin signaling pathways are both implicated in early brain development and the role of FEZ1 as a mediator of semaphorin signaling has been recently reported (Chua et al., 2021; Goshima et al., 2016; Mulligan and Cheyette, 2012).

Notably, DEGs in FEZ1-null hCOs were significantly enriched for genes associated with neurodevelopment and neurodegenerative disorders (Fig. 2F). FEZ1 abnormalities have been previously linked to Alzheimer’s and Parkinson’s disease as well as schizophrenia and ADHD (Butkevich et al., 2016; Sakae et al., 2008; Sumitomo et al., 2018; Sun et al., 2020). Additionally, genes associated with depression and intellectual disability were highly represented in FEZ1-null hCOs. Collectively, these results indicated FEZ1’s involvement in early human brain development and that loss of FEZ1 potentially triggers changes in biological pathways contributing to neurodevelopmental as well as neurodegenerative disorders.

### FEZ1-null hCOs contain a disproportionately larger population of HOPX^−^ oRG

Changes in expression of genes regulating neuronal differentiation were observed in FEZ1-null hCOs (Fig. 2E). This suggested *FEZ1* deletion may trigger alterations in developmental trajectories of neural progenitors. We performed single-cell RNA-seq (scRNA-seq) to interrogate for changes in sub-populations of cells in D28 hCOs when cortical lamination had already started (Camp et al., 2015; Lancaster et al., 2013; Qian et al., 2016). A total of 7800 and 9254 cells were analyzed from wildtype and FEZ1-null hCOs, respectively. Neural progenitors and neurons were identified and grouped according to known markers as previously reported (Fig. 3A and Fig S3A, Table S4) (Bhaduri et al., 2020; Nowakowski et al., 2017; Pollen et al., 2015). In total, 14 unique cell clusters could be identified, namely ventricular neuroepithelium/radial glia cells (vNE/vRG: *LIX1, NES, HMGA2, PAX6);* dividing vNE/vRG (*MKI67, LIX1, PAX6, ASPM);* truncated radial glia cells (tRG: *CRYAB, EGR1, HMGA2*); HOPX^+^ outer radial glia cells (HOPX^+^ oRG: *HOPX, BMP7, CLU, FEZF2*); HOPX^−^ outer radial glia cells (HOPX^−^ oRG: *FABP7, MOXD1, QKI),* which expressed oRG markers that are not represented in aforementioned HOPX^+^ oRG cluster (Bhaduri et al., 2020; Matsumoto et al., 2020; Nowakowski et al., 2017; Pollen et al., 2015); dividing oRG (*MIK67, HJURP, FABP7, CLU, FEZF2, HOPX);* choroid plexus (CP: *TTR, OTX2);* intermediate progenitor cells (IP: *EOMES, DCX, BEUROG1);* dividing IP (*EOMES*, *MKI67, NEUROG1*); new born neurons (*STMN2, DCX);* deep layer neurons (*STMN2, NEUROD6, BCL11B);* upper layer neurons (*POU3F2, BHLHE22);* interneurons (*DLX5, GAD2, DLX1);* and pericytes (*COL3A1,LUM).* Amongst neurons and their progenitor cells, both the proportion of cells expressing *FEZ1* as well as the level of *FEZ1* mRNA increased as progenitor cells sequentially develop into deep and upper layer neurons in wildtype hCOs, which agreed with results obtained by immunostaining (Fig. 3B, Fig. 1A).

**Figure 3.**
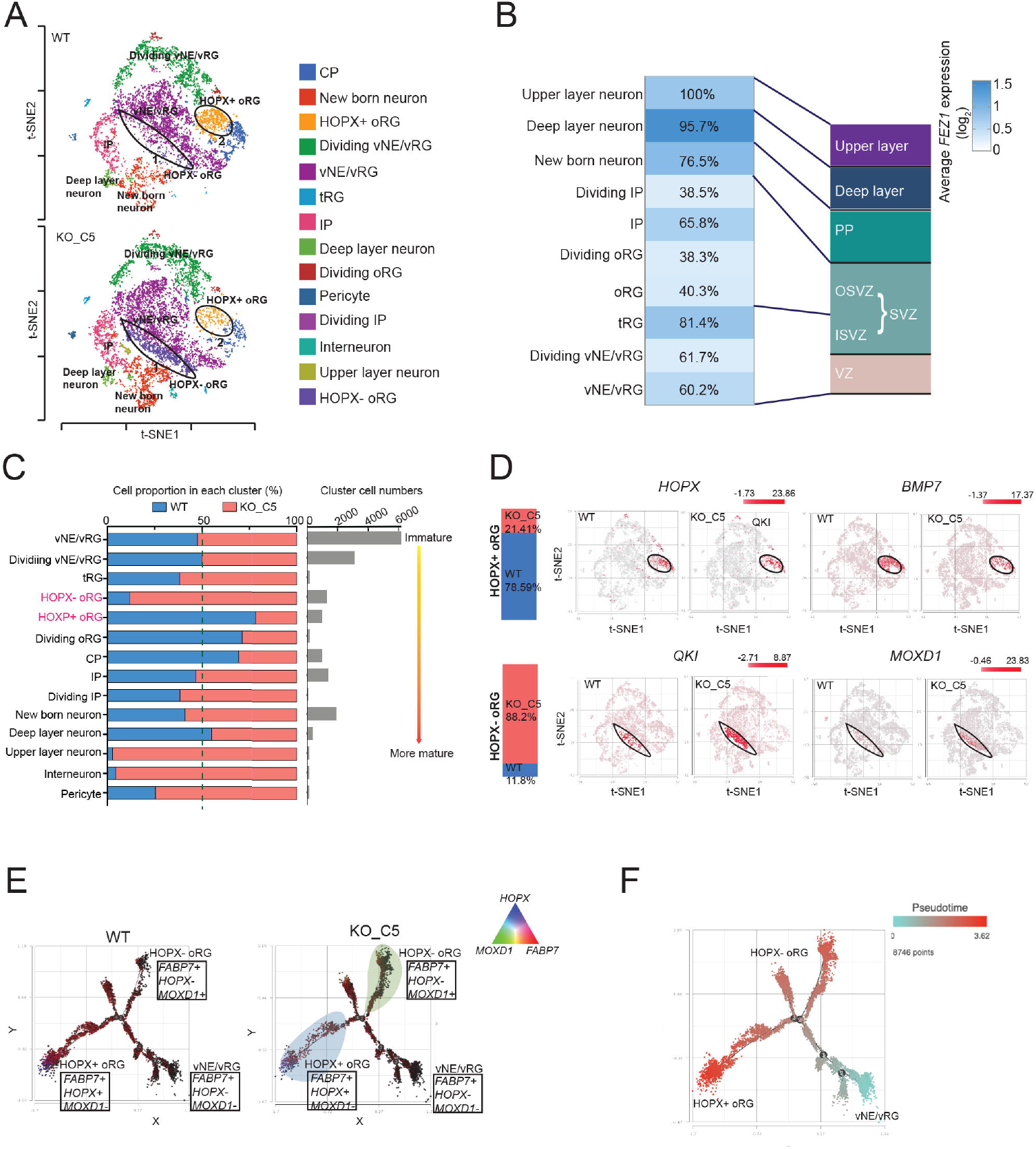
Single-cell RNA (scRNA) sequencing analyses uncovered alterations in development trajectories of outer radial glial (oRG) subpopulations in FEZ1-null hCOs. (A) t-SNE plots showing the distribution of various cell populations in D28 WT and FEZ1-null hCOs. Areas encircled with black ovals delineated HOPX^+^ oRG and HOPX^−^ oRG. vNE/vRG: ventricular neuroepithelium/ radial glia cells; tRG: truncated radial glia cells; HOPX^−^ oRG: HOPX^−^ outer radial glia cells; HOPX^+^ oRG: HOPX^+^ outer radial glia cells; CP: choroid plexus; IP: intermediate progenitor. (B) Relative expression levels of *FEZ1* mRNA in all classes of neuroprogenitor and newborn/immature neurons identified. Numbers in the boxes indicate percentages of cells expressing *FEZ1* in the corresponding cell clusters. (C) Proportion of cells in each cell cluster in WT versus FEZ1-null hCOs. Cell clusters are arranged in progressing levels of differentiation, beginning from vNE/vRG to upper layer neurons. Interneuron and pericyte clusters are indicated at the bottom of the chart. Total cell numbers identified in each cell cluster are plotted on the right panel. Clusters belonging to HOPX^+^ oRG and HOPX^−^ oRG showed the largest changes between the 2 hCO groups. (D) t-SNE plots showing expression of *HOPX* and *BMP7* in HOPX^+^ oRG, and *QKI* and *MOXD1* in HOPX^−^ oRG clusters of WT and FEZ1-null hCOs, respectively. (E) Trajectory inference analysis of three cell clusters (vNE/vRG, HOPX^+^ oRG and HOPX^−^ oRG) indicated that both HOPX^−^ oRG and HOPX^+^ oRG originated from vNE/vRG. In FEZ1-null hCOs, changes in developmental trajectories led to an increase in the HOPX^−^ oRG (shaded in green) and a decrease in the HOPX^+^ oRG (shaded in blue). (F) Pseudotime analysis indicated that HOPX^+^ oRGs were in a more mature state as compared to HOPX^−^ oRGs. (E) and (F) were analyzed using Partek Flow. VZ: ventricular zone; SVZ: sub-ventricular zone; ISVZ: inner sub-ventricular zone; OSVZ: outer sub-ventricular zone; PP: preplate.

Of the 14 unique cell clusters identified, changes in the proportion of HOPX^−^ oRG and HOPX^+^ oRG clusters were most apparent between FEZ1-null and wildtype hCOs (Fig. 3C). The percentage of HOPX^+^ oRG was dramatically reduced from 78.59% in wildtype hCOs to 21.41% in FEZ1-null hCOs. In comparison to this, the population of HOPX^−^ oRG increased substantially from 11.8% in wildtype HCOs to 88.2% in FEZ1 null hCOs (Fig. 3C). Both HOPX^+^ and HOPX^−^ oRG are present during cortical development, but their molecular signature remains incompletely defined (Matsumoto et al., 2020). We noticed that *FEZ1* expression was higher in HOPX^+^ oRG as compared to HOPX^−^ oRG in the pooled oRG populations, suggesting that *FEZ1* expression is required for, or at least, accompanies the development of HOPX^+^ oRG (Figure S3B). Closer examination of markers expressed by both clusters further showed that HOPX^+^ oRGs also expressed *BMP7,* markers characteristic of classical oRGs (Fig. 3D; Fig S3A; Table S4) (Bhaduri et al., 2020; Nowakowski et al., 2017; Pollen et al., 2015). In comparison to this, HOPX^−^ oRG did not express HOPX. Instead, they express *MOXD1, QKI* and *CLU,* which are also expressed in vRG and early-stage oRG (Bhaduri et al., 2020; Hayakawa-Yano et al., 2017; Pollen et al., 2015; Wang et al., 2011). Supporting these observations, the pseudotime value of HOPX^−^ oRG lies intermediate between vNE/vRG and HOPX^+^ oRG, suggesting HOPX^−^ oRG were at a more immature or earlier stage of development as compared to HOPX^+^ oRG (Haskell and LaMantia, 2005; Ji and Ji, 2016; Yu et al., 2019)(Fig. 3F). Both oRG populations initially originated from vNE/vRG as revealed by developmental trajectory analyses, but they subsequently diverged to form 2 discrete clusters (Fig. 3E). With loss of FEZ1, oRG developmental trajectory preferentially moved towards HOPX^−^ oRG expressing MOXD1 and QKI (Fig. 3E). This suggests that the generation of cell types downstream of these neuroprogenitors could be affected, which may alter subsequent brain development.

### Lamination abnormalities in FEZ1-null hCOs

In the primate brain, oRGs residing in SVZ are largely responsible for producing the diversity of cortical layer neurons during neocortex expansion (Penisson et al., 2019; Pollen et al., 2015). The developmental trajectory of oRG is of particular interest in neurodevelopmental disorders as it is unique to human and primate brain development and largely absent in mouse brains (Beattie and Hippenmeyer, 2017; Florio and Huttner, 2014; Molnar et al., 2019). Abnormalities associated with oRG subpopulation specification can contribute to cortical malformations that lead to alterations in neuronal connectivity (Juric-Sekhar and Hevner, 2019; Romero et al., 2018). A disproportionate increase in the number of HOPX^−^ oRG over classical HOPX^+^ oRG cells in FEZ1-null over wildtype hCOs is likely to alter subsequent generation of neuronal subtypes that would affect the eventual composition of cortical neurons as well as formation of cortical layers (Liu et al., 2017; Matsumoto et al., 2020; Nowakowski et al., 2017; Tabata et al., 2013).

To determine if cortical layer formation could be affected in the absence of FEZ1, D28 hCOs were stained for PAX6, TBR2 (IP) and CTIP2 (deep layer neurons) to mark the regions of VZ-ISVZ, OSVZ and PP, respectively (Lancaster and Knoblich, 2014; Pollen et al., 2015; Qian et al., 2016; Renner et al., 2017). Proper cortical lamination could be observed in wildtype hCOs, where VZ-ISVZ, OSVZ and PP layers were arranged concentrically from the centre to the outermost layers (Fig. 4A) (Bershteyn et al., 2017; Renner et al., 2017). In comparison, there was conspicuous overlapping of the TBR2^+^ OSVZ and CTIP2^+^ PP layers in FEZ1-null hCOs (Fig. 4A). In particular, the distribution of TBR2^+^ cells can be observed to encroached into the CTIP2^+^ layer. This was accompanied by a slight but significant decrease in percentage of TBR2^+^ cells in the OSVZ (27.2%±2,2% FEZ1-null hCOs; 34.7%±1.7%, wildtype hCOs) and a corresponding increase in the PP (28.9±2.7% in FEZ1-null hCOs; 22%±1.6% in wildtype hCOs) (Fig. 4B). Conversely, the percentage of CTIP2^+^ cells in the PP was reduced in FEZ1-null hCOs (38.5%±2.5% in FEZ1-null hCOs; 52.9%±2.5% in wildtype hCOs), which was accompanied by an increase in the OSVZ (27.4%±1.8% in FEZ1-null hCOs; 16.3%±1.1% in wildtype hCOs) (Fig. 4B). Defective lamination was also observed in hCOs generated from the FEZ1-null hiPSCs (Fig. S4A; Fig S2F). Thus, loss of FEZ1 indeed lead to lamination defects at the PP region during early corticogenesis in FEZ1-null hCOs (Fig. 4C).

**Figure 4.**
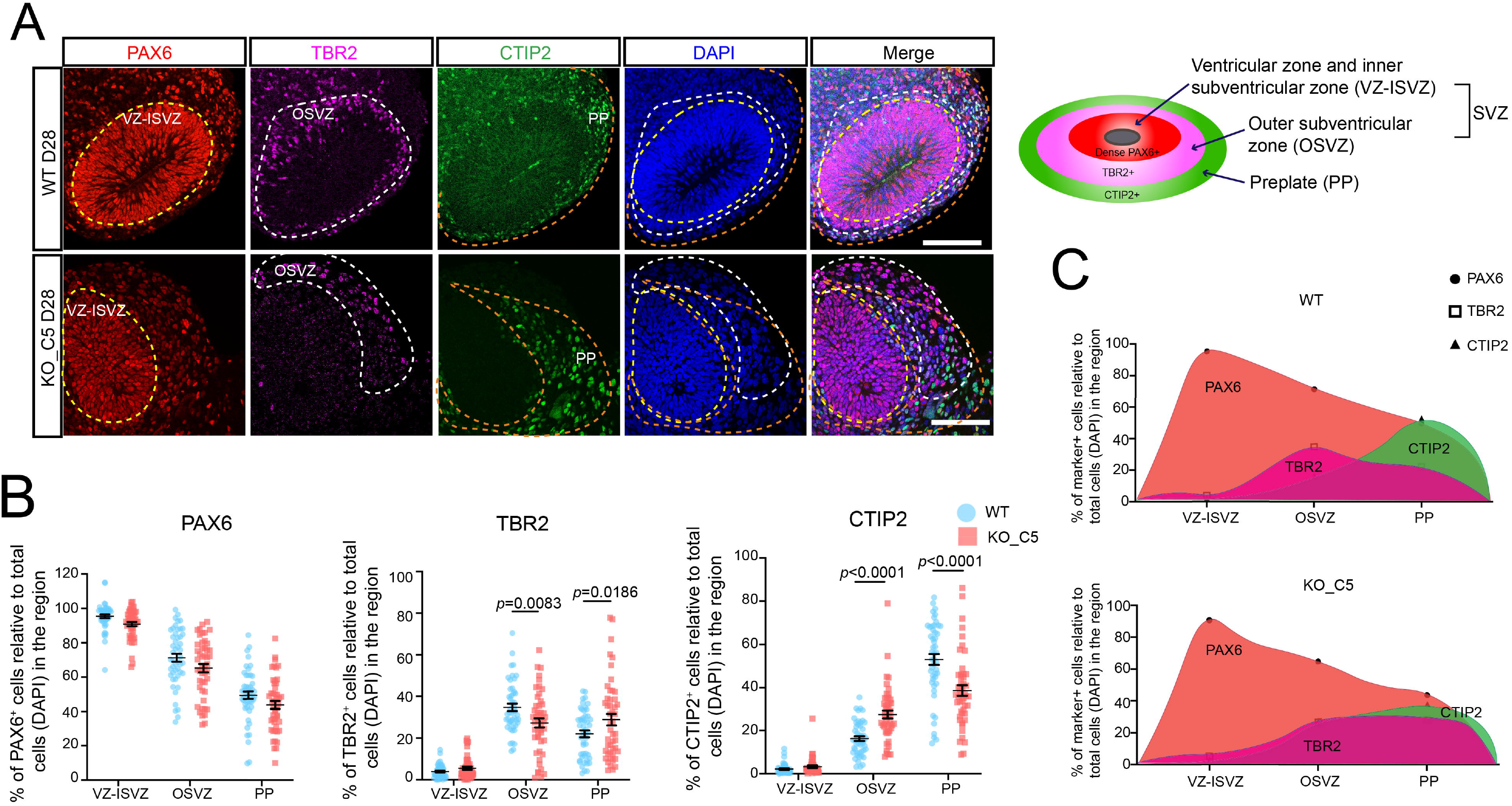
Cortical lamination defects were present in D28 FEZ1-null hCOs. (A) IF staining of PAX6, TBR2 and CTIP2 showed abnormal lamination in D28 FEZ1-null hCOs. Layering in WT hCOs was unaffected. Scale bar: 100 μm. (B) Quantification of PAX6^+^, TBR2^+^ and CTIP2^+^ cells in the VZ-ISVZ, OIVZ and PP in D28 WT and FEZ1-null hCOs. Values represent mean ± SEM (n= 4 independent organoid differentiations with total 9 organoids analyzed for WT and FEZ1-null group, respectively. Each data point represents one analyzed organoid region. At least 5 regions were analyzed within each organoid. Two-way ANOVA with Šidák’s multiple comparison was performed with the *p* values indicated). (C) Charts showing distribution of PAX6^+^, TBR2^+^ and CTIP2^+^ cells in WT versus FEZ1-null D28 hCOs based on quantification of IF images as shown in (A). In WT hCOs, TBR2^+^ and CTIP2^+^ regions sequentially expand from OSVZ to PP. However, delineation of the 2 layers was perturbed in FEZ1-null hCOs and both cell types appeared to intermingle with each other.

Next, we examined if cortical layering abnormalities could be captured in our scRNA-seq dataset. Based on the quantification of cell population via scRNA-seq, a decrease in the number of CTIP2^+^ deep layer neurons in FEZ1-null hCOs was detected (1.9% in FEZ1-null hCOs; 2.3% in wildtype hCOs) (Fig. 3C; Fig. S4B). Conversely, the pool of TBR2^+^ IPs was expanded in FEZ1-null hCOs versus wildtype hCOs (8.6% in FEZ1-null hCOs; 7.6% in wildtype hCOs) (Fig. 3C; Fig S4B). Collectively, these results indicated that production of IP and the specification of cortical lamination, in particular of deep layer neurons are dysregulated in FEZ1-null hCOs.

### HOPX^−^ oRG exhibit higher levels of actin dynamics

To gain further insight into how the shift towards HOPX^−^ oRG causes lamination defects, we examined which genes were differentially expressed in HOPX^−^ versus HOPX^+^ oRG cells. DEGs between the two populations of oRG were extracted from wildtype and FEZ1-null hCOs and visualized using the BBrowser (Venice algorithm (FDR<0.05)) (Fig. 5A, Table S5) (Korthauer et al., 2016; Le et al., 2020; Wang et al., 2019). Genes that were most significantly upregulated in the HOPX^+^ oRG cluster included *FEZF2, NFIA, SYT1, STMN1, JGA1, BCL2* and *HOPX* (Fig. 5A; Table S5). In contrast to this, most significantly upregulated genes in HOPX^−^ oRG included *PAX6, FOXG1, FABP5, QKI* and *MOXD1* (Fig. 5A; Table S5). DEGs in HOPX^−^ oRG were enriched for biological processes related to cerebral cortex development, neuronal differentiation, axogenesis, microtubule cytoskeleton organization, supporting the notion that the increased in population of HOPX^−^ oRG accompanying FEZ1 loss can affect early brain development (Fig. 5B; Table S6). Analyses of DEGs against KEGG curated pathways further highlighted changes in expression of genes involved in MAPK signalling, regulation of actin cytoskeleton, cell adhesion molecules, tight junction and focal adhesion (Fig. S5A, Table S7). Of note, significant alterations in expression of genes involved in cell adhesion and actin cytoskeleton regulation (Figure S5B) (Table S7), such as ACTB (log^2^FC = −0.61, FDR = 9.25e-77), CDH2 (log^2^FC = 0.393; FDR = 5.20e-24), NCAM1 (log^2^FC = 0.22; FDR = 6.9e-12), CNTNAP2 (log^2^FC = −2.37; FDR<1.89E-294), CNTNAP3B (log^2^FC = −0.25; FDR = 4.17E-22), PTK2 (log^2^FC = −0.147; FDR = 4.76E-8), MAPK10 (log^2^FC = −0.5; FDR = 3.71E-50), MAPK3 (log^2^FC = −0.2; FDR = 2.17E-19) were detected in HOPX^−^ oRG (Table S5). Both signaling pathways are crucial for neuronal morphogenesis and cell migration during corticogenesis (Huang et al., 2020; Solecki et al., 2009; Vicente-Manzanares et al., 2009). Additionally, significant downregulation of TMSB4X (log^2^FC = −0.398; FDR = 1.85E-15), PFN1 (log^2^FC = −0.212; FDR = 7.35E-09), PFN2 (log^2^FC = −0.341; FDR = 4.31E-20) and ARPC2 (log^2^FC = −0.162; FDR = 4.45E-6) were also observed in HOPX^−^ oRG (Fig. 5C, Table S5). These molecules are known to regulate F-actin polymerization, which is vital in modulating cell migration and neurite outgrowth of neural progenitors (Fig. 5D) (Huang et al., 2020; Padmanabhan et al., 2020; Schaks et al., 2019). Therefore, we examined whether alterations in actin dynamics were present in neuronal progenitor cells following loss of FEZ1.

**Figure 5.**
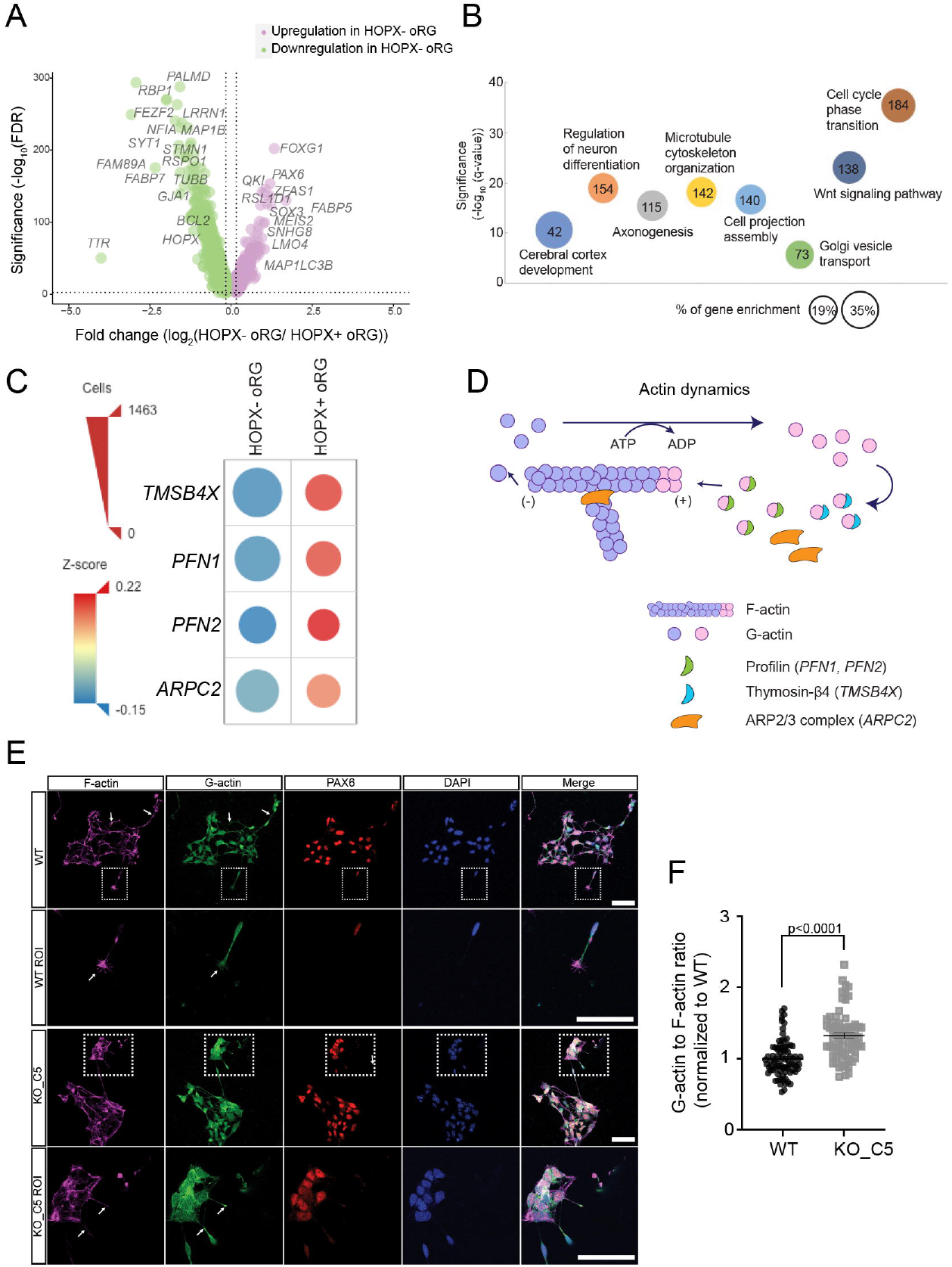
HOPX^−^ oRG population exhibited higher levels of actin dynamics. (A) Volcano plot of DEGs in HOPX^−^ oRGs compared with HOPX^+^ oRGs. DEGs of interest are indicated in the plot. (B) GO enrichment analyses (by Metascape) of DEGs in HOPX^−^ oRGs as compared to HOPX^+^ oRGs. Size and number of the circles represent the percentage and number of DEGs involved in each term, respectively. (C) Dot plot showing expression levels of DEGs involved in actin polymerization in HOPX^−^ oRGs and HOPX^+^ oRGs. Affected genes were downregulated in HOPX^−^ oRGs. Expression levels are color-coded according to their Z-scores. The diameter of each dot correlates with the number of cells (as scaled by triangular red legend) for the corresponding gene. Analysis was done using Bioturing. (D) Diagram showing the regulation of actin dynamics. Polymerized F-actin is dissembled into monomer G-actin at the minus (-) end while G-actin is incorporated at the plus (+) end of F-actin. Profilin 1 and 2 (*PFN1, PFN2),* Thymosin-β4 (*TMSB4X),* ARP2/3 complex (*ARPC2*) participate in assembling monomer G-actin to polymerized F-actin. (E) IF staining of F- and G-actin in WT and FEZ1-null PAX6^+^ neural progenitors dissociated from D28 hCOs. Arrows in ROIs shows enrichment of G-actin at neurite tips in FEZ1-null neural progenitors as compared with WT. Scale bar: 100 μm. (F) Quantification of G- to F-actin ratio in neural progenitors dissociated from D28 hCOs from WT and FEZ1-null hCOs. Values represent mean ± SEM (n=3 independent organoid differentiations for WT and FEZ1-null group, respectively. Each data point represents measurements from one image, at least 10 images were analyzed within each replicate. Unpaired t-test were performed to determine the two-tailed *p* value as indicated).

Changes in actin polymerization can be assessed by measuring the ratio of depolymerized (globular (G)-actin) to polymerized (filamentous (F)-actin) actin (Huang et al., 2020). We dissociated cells from D28 hCOs, plated them on coverslips and immunostained for G- and F-actin together with PAX6, a neuroprogenitor cell marker that has been reported to localized to RG cells (Fig. 5E) (Gotz et al., 1998; Hansen et al., 2010; Matsumoto et al., 2020). The G- to F-actin ratio in PAX6^+^ RG cells was calculated by measuring total fluorescent intensities on each channel, respectively (Fig. 5E). We postulated that any changes in G- to F-actin ratio in PAX6^+^ cells between FEZ1-null and wildtype hCOs is likely attributed to differences in the relative proportion of HOPX^+^ and HOPX^−^ oRG populations. PAX6^+^ RG cells from FEZ1-null hCOs indeed exhibited 33.6%±5.4% higher G- to F-actin ratio as compared to those from wildtype hCOs (Fig. 5F). G-actin also appeared to be more enriched at the tips of processes in cells from FEZ1-null hCOs (Fig. 5E). Thus, the larger proportion of HOPX^−^ oRGs exhibiting higher levels of actin dynamics can account for layering abnormalities in FEZ-1 null hCOs.

### Ectopic localization and atypical cell migration of HOPX^−^ oRG in FEZ1-null hCOs during corticogenesis

Migration of oRG to OSVZ-PP during early cortical brain development is crucial for proper localization and expansion of the cortical layers (Marin et al., 2010; Taverna et al., 2014). Migration abnormalities give rise to cortical developmental disorders (Juric-Sekhar and Hevner, 2019; Silva et al., 2019). We next proceeded to investigate how the disproportionate amount of oRG with higher actin dynamics contributes to cortical layer formation abnormalities in FEZ1-null hCOs.

There is currently no reliable marker to distinguish between HOPX^+^ versus HOPX^−^ oRG. Since *QKI* was highly expressed in HOPX^−^ but not in HOPX^+^ oRG (Figure S3A), we employed it as a putative marker to examine the distribution of the former population of cells in D28 hCOs. In wildtype D28 hCOs, QKI^+^ oRG were predominantly found in the inner SVZ region (ISVZ) and rarely at the outer region of ISVZ (including the OSVZ and PP region, henceforth collectively indicated as OSVZ-PP) (Fig 6A). This supports its expression in vRG and oRG at this stage of development (Hayakawa-Yano et al., 2017; Pollen et al., 2015). Remarkably, OSVZ-PP in D28 FEZ1-null hCOs contained significantly more of QKI^+^ oRG as compared to wildtype hCOs (Fig. 6A). Similar results were observed using MOXD1, which is also highly expressed in HOPX^−^ oRG. More MOXD1^+^ cells were found in the OSVZ-PP of FEZ1-null hCOs as compared to wildtype hCOs (Fig. S6A).

**Figure 6.**
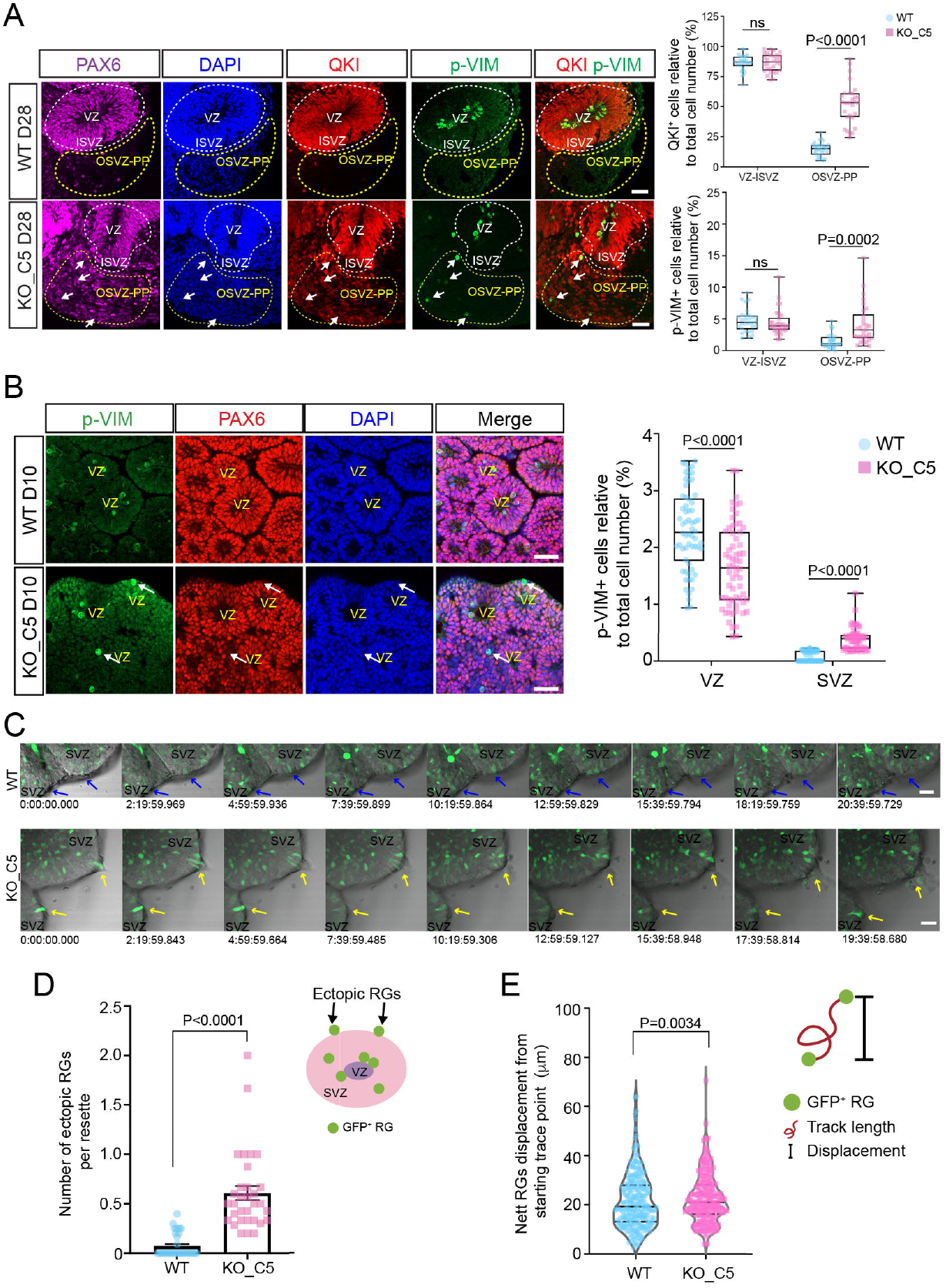
Ectopic localization of HOPX^−^ oRG in FEZ1-null hCOs. (A) D28 WT and FEZ1-null hCOs were stained for PAX6 (RG marker), QKI (vRG and HOPX-oRG marker) and p-VIM (mitotic RG marker). In WT hCOs, QKI^+^ RGs and mitotic RGs were largely confined within the ISVZ. In FEZ1-null hCOs, the populations of both cell types increased dramatically in the OSVZ-PP region. White arrows indicate pVIM^+^ mitotic RGs. Quantification of QKI^+^ and p-VIM^+^ cells are shown on the right. Scale bars: 50 μm. (B) Ectopic mitotic RG (p-VIM^+^) could be observed at the outer surface of rosettes in D10 FEZ1-null hCOs (arrows). Quantification of p-VIM^+^ cells are shown on the right. Scale bars: 100 μm. For (A) and (B), values represent mean ± SEM (n=3 independent organoid differentiations with total 8 organoids analyzed for WT and FEZ1-null groups, respectively. Each data point represents one analyzed region of an organoid; at least 5 different regions were analyzed from one organoid. Unpaired t test with Welch’s correction was performed, with two-tailed *p* values as indicated). (C) Time lapse imaging in D10 hCOs infected with GFP-expressing AAVs showed abnormal migration of GFP^+^ RG in FEZ1-null hCOs as compared to WT hCOs. Arrows indicate labelled cells at the edge of SVZ. Cells in WT hCO were confined within each rosette (VZ/SVZ region) while undergoing proliferation or relocation (blue arrows). In contrast, some RG cells in FEZ1-null hCOs appear atypical migration behaviour by migrating out of the rosette (SVZ edge region, yellow arrow). Corresponding videos can be found in supplement video 1 (D10 WT hCO migration) and video 2 (D10 FEZ1-null hCO migration). Scale bars: 50 μm. (D) Illustration and quantification of atypical cells with abnormal migration observed during time lapse recording. Values represent mean ± SEM (n=3 independent organoid differentiations for WT and FEZ1-null group, respectively. Each data point represents one analyzed region in an organoid. At least 10 different regions were analyzed within one replicate. Unpaired t-test with Welch’s correction was performed with two-tailed *p* values as indicated). (E) Quantification of nett cell displacement. Nett displacement of GFP^+^ RGs in FEZ1-null hCOs was higher as compared to WT hCOs. Values represent mean ± SEM (n=3 independent organoid differentiations, each data point represents one traced cell that could be consecutively traced for more than 7 hours. Unpaired t-test with Welch’s correction was performed with two-tailed *p* values as indicated). VZ: ventricular zone; SVZ: sub-ventricular zone; ISVZ: inner sub-ventricular zone; OSVZ-PP: outer sub-ventricular zone-preplate.

We noticed that ectopic localization of QKI^+^ oRG could already be seen in D10 FEZ1-null hCOs (Fig. S6B). In wildtype hCOs, pVIM^+^ mitotic RG were typically found close to VZs and rarely at the edge of each rosette at both D10 and D28 wildtype hCOs (Fig. 6A-B). However, the ectopic mitotic RG were often detected at the outer edge of rosettes at the OSVZ regions in D10 FEZ-1-null hCOs, with a concomitant reduction in the number of VZ-associated mitotic RGs (Fig. 6B). This population of ectopic pVIM^+^ RGs in the OSVZ-PP region persisted as the mutant hCOs developed (Figure 6A, D28), therefore indicating loss of FEZ1 not only resulted in an increase and persistence of HOPX^−^ oRGs, but also in their ectopic localization to the OSVZ-PP regions of the hCOs.

As our previous scRNAseq data highlighted differences actin dynamics between HOPX^−^ and HOPX^+^ oRG, we further examined whether the ectopically localized oRG population exhibited abnormal cell migratory behaviour by infecting D10 FEZ1-null and wildtype hCOs with a low titre of GFP-expressing AAVs to sparsely labelled cells. We tracked the migration of GFP^+^ cells over a 20 h period by time-lapsed imaging. Most cells present at this stage are presumed to be dominantly PAX6^+^ RG (Gotz et al., 1998; Hansen et al., 2010; Matsumoto et al., 2020). In agreement with the presence of an expanded population of HOPX^−^ oRG cells with higher actin dynamics with FEZ1 loss, a higher proportion of migrating GFP-labelled cells could be observed to move to the outer edge of rosettes in FEZ1-null hCOs (yellow arrow) versus wildtype hCOs (Fig. 6C and D). Interestingly, more GFP^+^ cells in mutant hCOs could be consecutively tracked for more than 7 hours, which suggested that they exhibited more persistent migratory behavior than those in wildtype hCOs (Fig. S6C; Supplementary Videos 1 (WT) and 2 (FEZ1-null)). Additionally, these cells in FEZ1-null hCOs showed greater displacement (Fig. 6E) even though track length and migration speed did not show significant difference as compared to WT cells (Fig. S6D and E). Furthermore, these cells appear to remain at the periphery, in line with the increased number of ectopic oRG in FEZ1-null hCOs.

Previous studies indicated HOPX^−^ oRG possessed both self-renewal capability while were able to further differentiate into IP and other neuronal cell types (Matsumoto et al., 2020). To further investigate how ectopic HOPX^−^ oRG could affect cortical layering, we tested the self-renewal capability of these cells by pulse labelling the organoids with 5-ethynyl-2’-deoxyuridine (EdU) on D23 and harvesting them D28, when HOPX^−^ oRG population can be detected in OSVZ region (Figure 6B). We defined the dense PAX6^+^ region as VZ-ISVZ and the region that was strongly labelled for TUJ1 or DCX surrounding the VZ-ISVZ as OSVZ-PP (Li et al., 2017; Qian et al., 2016; Renner et al., 2017) (Fig. 7, A and B). In contrast to wildtype hCOs, more EdU^+^ post-mitotic cells were observed at the outer boundaries OSVZ-PP in FEZ1-null hCOs. A portion of these cells also stained positive for PAX6 (Fig. 7C, arrows). We next determined where cell proliferation occurs by staining for Ki67. Increased amount of mislocalized mitotic (Ki67^+^) cells in OSVZ-PP of FEZ1-null hCOs were observed (Fig. S7, A and B). A proportion of Ki67^+^ cells were also PAX6 (Fig. S7C, arrow). Together with the earlier results showing localization of ectopic HOPX^−^ oRG, these data indicated that abnormal migration and subsequent division of mislocalized oRG eventually resulted in abnormalities in cortical layer formation in FEZ1-null hCOs.

**Figure 7.**
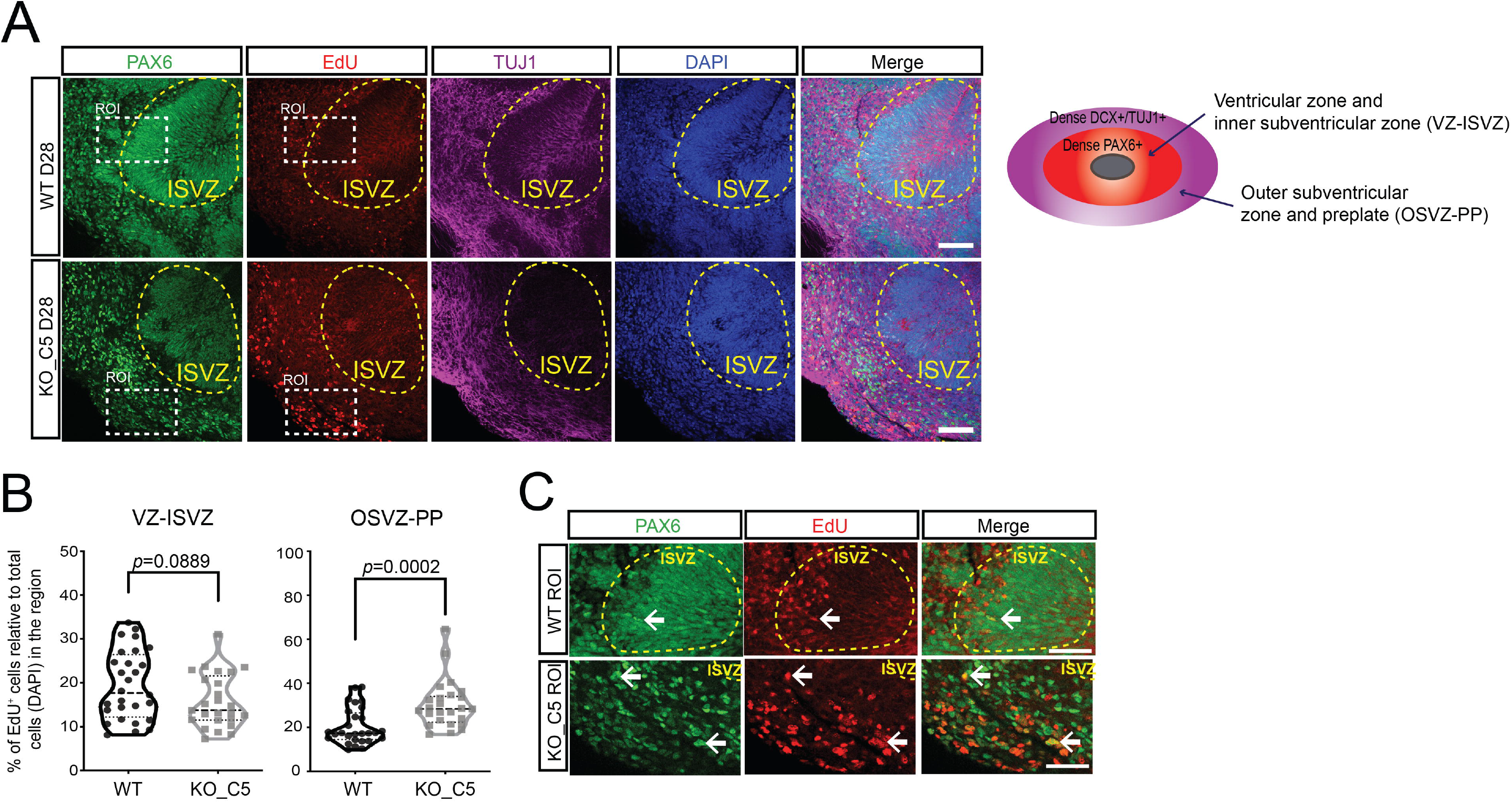
Proposed mechanism of early lamination defect in FEZ1-null hCOs. (A) Lineage tracing of postmitotic daughter cells on D28 hCO by EdU pulse labelling assay for both WT and FEZ1-null group with co-staining with PAX6 and TUJ1. More daughter cells were located at the OSVZ region in FEZ1-null hCOs. Scale bars: 100 μm. (B) Quantification of EdU^+^ post-mitotic cells in different regions. Values are represented as violin plots with median and quartiles indicated as dashed and dotted lines, respectively. (n= 3 independent organoid differentiations with total 6 organoids analyzed for WT and FEZ1-null group, respectively. Each data point represents value from one analyzed organoid region. At least 4 regions were analyzed for each organoid. Unpaired t-test with Mann-Whitney test was applied to determine the indicated two-tailed *p* values). (C) ROI in (B) showing that a proportion of EdU^+^ cells are PAX6^+^ RGs (white arrow). Scale bars: 50 μm.

## Discussion

Using hCOs as a model of human brain development, we demonstrated that *FEZ1* expression was switched on early during cortical development as hESCs differentiated into neuroprogenitor cell types. Its levels continued to increase in major subsets of neuroprogenitors and was almost universally expressed in immature neurons. Deleting *FEZ1* significantly altered the expression of other genes involved in neuronal development and synaptic function. Remarkably, FEZ1 deficiency disrupted the ratio of HOPX^+^ and HOPX^−^ oRG, with HOPX^−^ oRG being substantially increased and HOPX^+^ oRG being decreased in FEZ1-null hCOs. FEZ1-null HOPX^−^ oRG displayed differential expression of genes that regulate actin dynamics. Supporting the expansion of this oRG subtype, neuroprogenitor cells in FEZ1-null organoids exhibited increased levels of actin dynamics and had a higher tendency to migrate to the peripheral edge of the organoids. This observation was suggestive of altered cell migration patterns in FEZ1-null hCOs, which were supported by greater inter-mixing of TBR2^+^ intermediate progenitor cells with CTIP2^+^ deep layer neurons in FEZ1-null hCOs, indicative of abnormalities of early cortical layer formation. Collectively, these results highlight a hitherto unknown role of FEZ1 during the early stages of brain development.

Although previous reports highlighted the involvement of FEZ1 in neuronal and synaptic development, its involvement in the earlier stages of brain development has never been fully explored (Bloom and Horvitz, 1997; Butkevich et al., 2016; Chua et al., 2021; Miyoshi et al., 2003). Detection of *FEZ1* transcripts early in rodent brain development and the association of *FEZ1* mutations in human neurodevelopmental disorders indicate the possibility of its involvement at even earlier stages of brain development (Razar et al., 2022). Indeed, *FEZ1* mRNA and protein were already present as early as D10 in hCOs, which is equivalent to a 4-5 weeks-old human embryo and an E11 mouse embryo (Bystron et al., 2006; Camp et al., 2015; Fujita et al., 2004). Moreover, scRNA-seq and confocal microscopy analyses further localized *FEZ1* expression to distinct subsets of neuroprogenitor cells, including NE, vRG, oRG and, later, in newborn as well as immature neurons.

Early in the course of neurogenesis, progressive expansion and differentiation of NE to vRG, oRG and IP are responsible for producing the vast diversity of cortical neurons and subsequent organized expansion of cortical layers (Gertz et al., 2014; Greig et al., 2013; Nowakowski et al., 2017; Pollen et al., 2015). oRGs play a pivotal role during the development of the human cortex by producing intermediate progenitor cells (IP), which in turn give rise to neurons and various types of glial cells, contributing to cortical expansion and lamination, and the eventual formation of different cerebral regions (Molnar et al., 2019). oRGs in the OSVZ can be further distinguished into 2 subtypes by the selective expression of *HOPX.* HOPX^+^ oRG form the larger proportion that are present in human OSVZ (Matsumoto et al., 2020; Pollen et al., 2015). Loss of *FEZ1* appears to mainly affect oRG. A substantial increase in the proportion of HOPX^−^ oRG, accompanied by a corresponding decrease in HOPX^+^ oRG was detected in FEZ1-null hCOs as compared to wildtype hCOs. HOPX^+^ oRG and HOPX^−^ oRG show differences in their ability to generate TBR2^+^ IP (Matsumoto et al., 2020). The shift in both quantity and proportion of both oRG types was accompanied by a disproportionate increase in TBR2^+^ IP and reduction in CTIP2^+^ deep layer neurons and encroachment of IP into the deep layer neuron layer. In agreement with this, our scRNAseq data uncovered an increase in the TBR2^+^ IP population in FEZ1-null hCOs (Figure S4). Moreover, upregulation of EOMES/TBR2 and INSM1 (upstream of TBR2 signaling), and downregulation of ZNFPM2/FOG2 (the layer 6 marker and downstream of TBR2 signaling) were observed in FEZ1-null hCOs compared with WT (Table S5), indicating aberrant IP production and transition to deep layer neurons in FEZ1-null hCOs (Cánovas et al., 2015; Mihalas et al., 2016; Wiegreffe et al., 2015; Woodworth et al., 2016). These results are reminiscent of abnormalities observed in *Bcl11a/CTIP1-deficient* mice, where an significant bias in specification and generation of subcerebral projection neurons and impaired neuronal radial migration was observed (Wiegreffe et al., 2015; Woodworth et al., 2016). Nevertheless, the significance of this relationship is at the moment unclear and is an important question for further investigation.

Apart from abnormalities in generation of neuroprogenitor subtypes, alterations in RG migration can also affect cortical formation (Hatten, 1999; Rakic, 1988). Mispositioning of oRGs as a result of dysregulated mammalian target of rapamycin (mTOR) signalling has been identified to affect oRG migration by changing their actin cytoskeleton through Rho-GTPase and CDC42 (Andrews et al., 2020). Although HOPX^+^ and HOPX^−^ oRG are both implicated in cortical expansion, differences between both cell types remain unclear (Liu et al., 2017; Matsumoto et al., 2020; Nowakowski et al., 2017; Tabata et al., 2013). Nevertheless, differences in actin cytoskeleton regulation has been suggested to contribute to differences in migration behaviour between the HOPX^+^ and HOPX^−^ oRG (Andrews et al., 2020; Tabata et al., 2009). Our examination of DEGs in HOPX^−^ oRG versus HOPX^+^ oRG also found changes in the expression of genes regulating actin dynamics, which generate protrusive and contractile motion involved in cell migration, especially during soma translocation (Schaks et al., 2019; Steinecke et al., 2014). The dynamics of stabilized and destabilized F-actin at the leading process tip of the cell provided pulling forces to drive nucleus moving, therefore to control cell migration and position (Steinecke et al., 2014).

While FEZ1’s function has been closely linked to microtubule-base intracellular transport, its association with the actin cytoskeleton has also been previously reported. FEZ1 was copurified with F-actin, through which it recruits DISC1 (Miyoshi et al., 2003). DISC1 has been shown to modulate neuronal migration through its interaction with actin cytoskeleton related proteins that either facilitate or inhibit neural migration, such as NDEL1, ACTB, AKT1 and GIRDIN (Enomoto et al., 2009; John et al., 2019; Steinecke et al., 2014). Hence, FEZ1 could exert its effect on the actin cytoskeleton and cell migration via its interaction with DISC1 (Miyoshi et al., 2003; Steinecke et al., 2014). Additionally, FEZ1 colocalized with actin in the tips of growing processes in oligodendroglia progenitor cells (Chen et al., 2017). Knocking down *FEZ1* expression in these cells significantly reduced the growth of these processes, suggesting that FEZ1 could be involved in regulating cell migration behaviour. Thus, together with the expanded population, higher levels of actin dynamics in HOPX^−^ oRG neuroprogenitors can account for the greater anomalous migration of neuroprogenitors towards the outer periphery of organoids observed in FEZ1-null hCOs.

In conclusion, our study uncovered the importance of FEZ1 during early forebrain development in terms of neurogenesis, neural specification and cortical brain deep layer stratification. These findings potentially contribute towards the understanding of neural developmental pathologies underlying neuropsychiatric disorders, therefore paving the way for potential treatment approaches.

## Acknowledgements

The authors thank Chua Jie Yin and Tang Jia Ying for producing lentiviruses used to generate FEZ1-null hPSCs. We thank Rafhanah Banu Bte Abdul Razar and Abigail Ruth Reyes Guillermo for helping with off-target checking of FEZ1-null hESC H1 and hiPSC. Confocal microscopy was supported by NUS The N.1 Institute for Health. We would also like to thank NUS Tissue Engineering Programme (NUSTEP) for supporting the stem cell and organoid culture. This work was supported by funding from the National University of Singapore to JJEC.

## Supplementary figures

**Figure S1.**
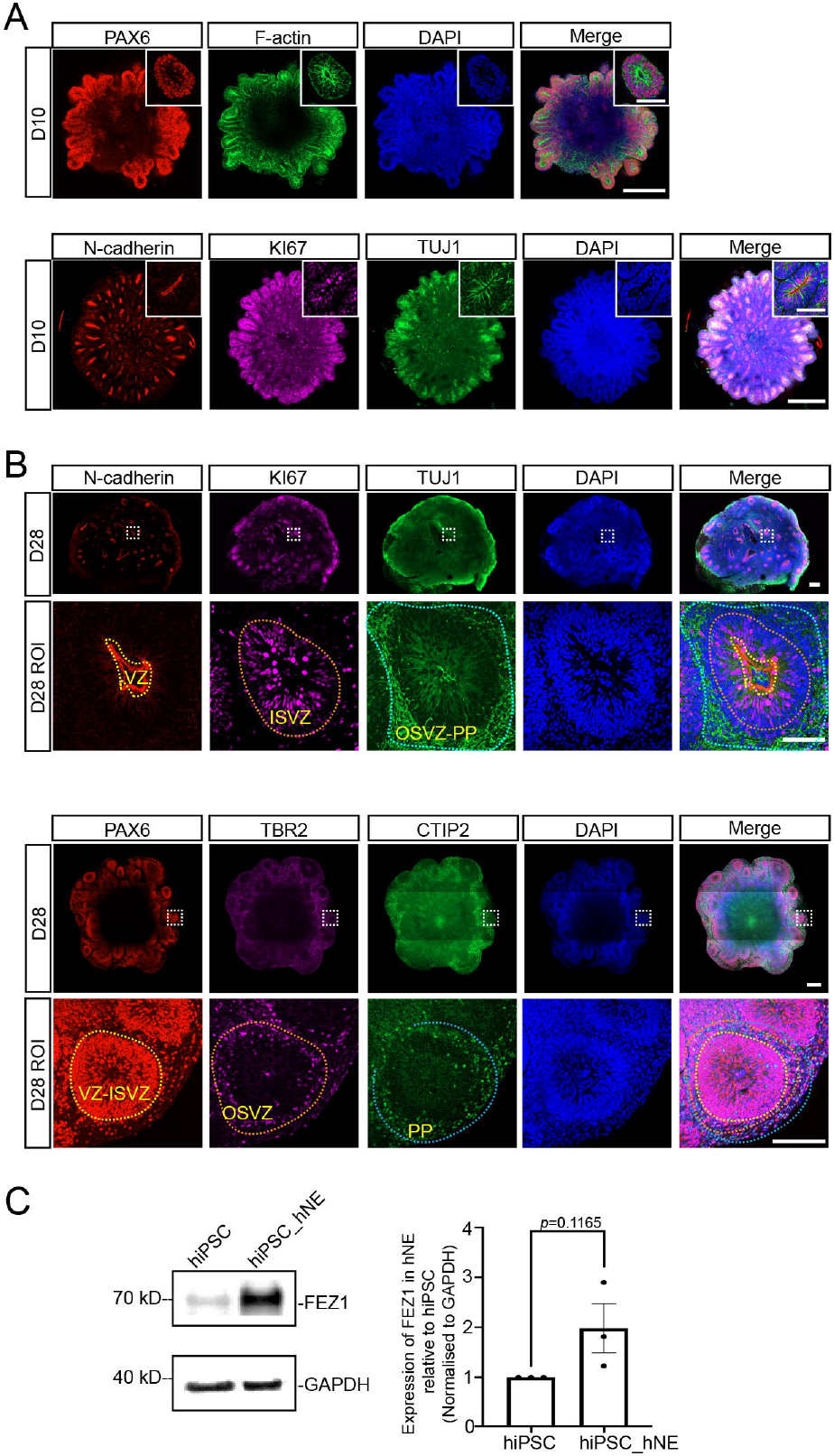
Characterization of hESC derived hCOs and FEZ1 expression in hiPSC derived neuroepithelial cells (NE) in 2D monolayer cultures. (A-C) PAX6 (NE/RG marker), F-actin, N-cadherin (apical/VZ marker), KI67 (proliferation marker) and TUJ1 (neuronal marker) staining in D10 (A) and (B), D28 (C) hCOs generated from H1 hESC confirmed hCOs established apical-basal polarity during development. Mitotic cells were predominately located near VZ region. (D) IF staining of D28 hCOs with PAX6 (NE/RG marker), TBR2 (IP marker) and CTIP2 (deep layer marker) validated cortical layering structure of VZ-ISVZ (PAX6), OSVZ (TBR2) to PP (CTIP2) established in D28 hCOs. Higher magnification of representative single rosette is shown in the upper right corner. Region of interest (ROI) as labelled in white dotted line square were displayed under corresponding images. Colored dotted lines indicated distinct layer boundaries in developing hCOs. Scale bars: 250 μm (A), (B), (C) and (D); 50 μm in upper right images (A) and (B). Immunoblot (E) and quantification (F) of FEZ1 protein expression in hiPSC and hiPSC derived neuroepithelial cells (NE) as a 2D monolayer. Values represent mean ± SEM. (n=3 independent hiPSC to hNE differentiations, unpaired t test determined the two-tailed *p* value as indicated). VZ: ventricular zone; ISVZ: inner sub-ventricular zone; OSVZ: outer sub-ventricular zone; PP: preplate.

**Figure S2.**
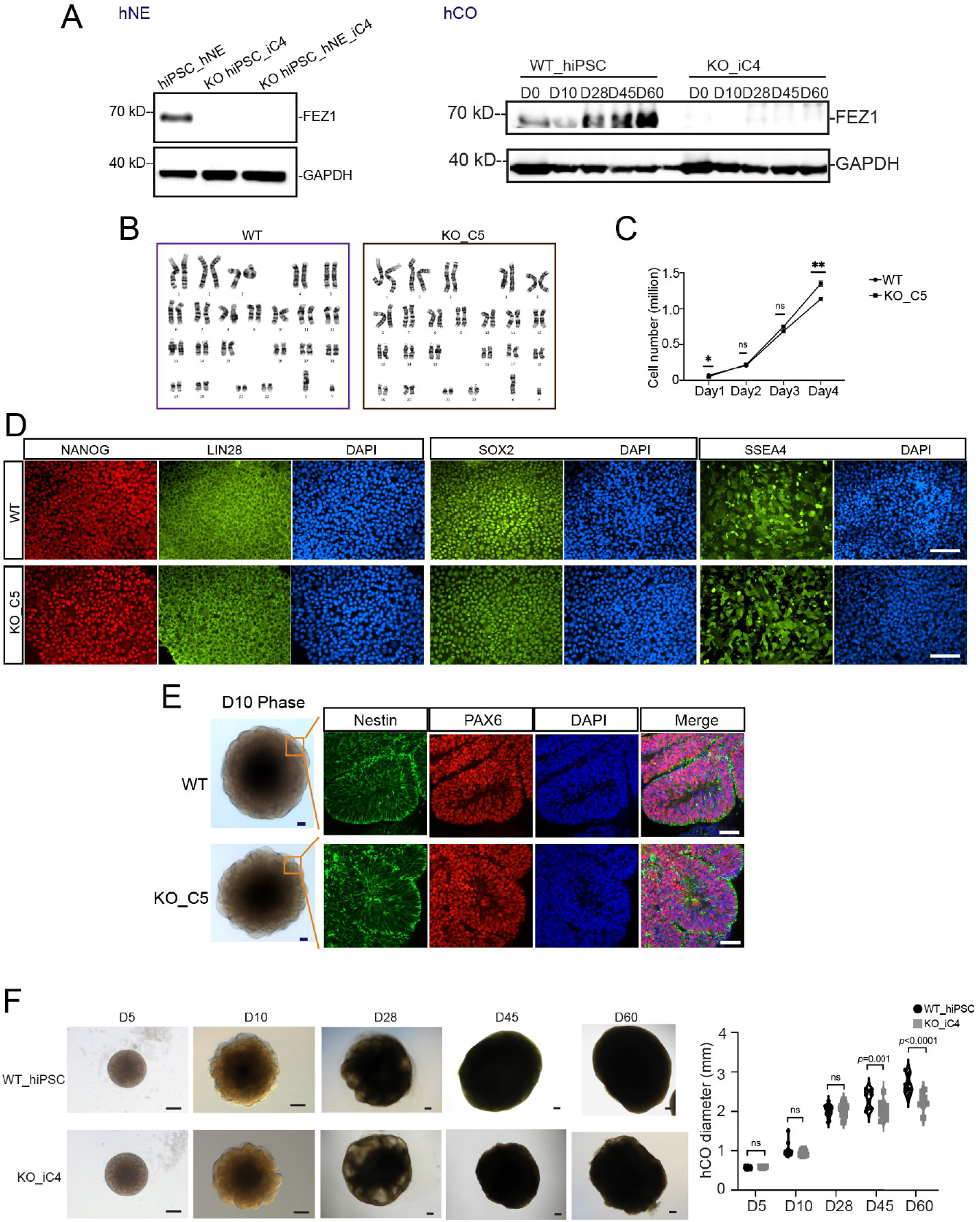
*Characterization of FEZ1-null hPSC and hCOs* derived from hiPSC *with loss of FEZ1.* (A) Immunoblot of FEZ1 expression in neuroepithelial cells (NE) and developing hCOs confirmed successful ablation of FEZ1 expression in FEZ1-null hiPSC_iC4 clone. (B) Karyotyping of wildtype (WT) H1 and FEZ1 KO H1 revealed no abnormality in both cells. 23 WT cells and 20 FEZ1 KO cells were analyzed. (C) Cell number count from Day 1 to Day 4 of cultured WT and FEZ1-null hESC H1, respectively. No obvious cell number difference was found during stem cell growth. Values represent mean ± SEM (n=3 independent experiments, two-way ANOVA with Šidák’s multiple comparison is used to determine the indicated *p* values) (D) IF staining of pluripotent makers in both WT H1 and FEZ1-null hESC H1 suggested FEZ1-null H1 maintained stem cell pluripotency. Scale bar: 50 μm. (E) Phase contrast images and IF staining of NESTIN and PAX6 in D10 WT and FEZ1-null hCOs. Neural rosettes were able to form in both WT and FEZ1-null organoids. Scale bar: 50 μm. (F) hCOs generated from WT hiPSC line and FEZ1-null hiPSC_iC4 clone showed no gross differences in morphology and size up to 28 days culture. Scale bar: 200 μm. (n=3 independent organoid differentiations up to 28 days and 2 independent organoid differentiations from 28 days to 60 days for WT and FEZ1-null group, respectively. Each data point represents value from one organoid, at least 4 organoids were measured in each replicate, two-way ANOVA with Šidák’s multiple comparisons, ns: not statistically significant).

**Figure S3.**
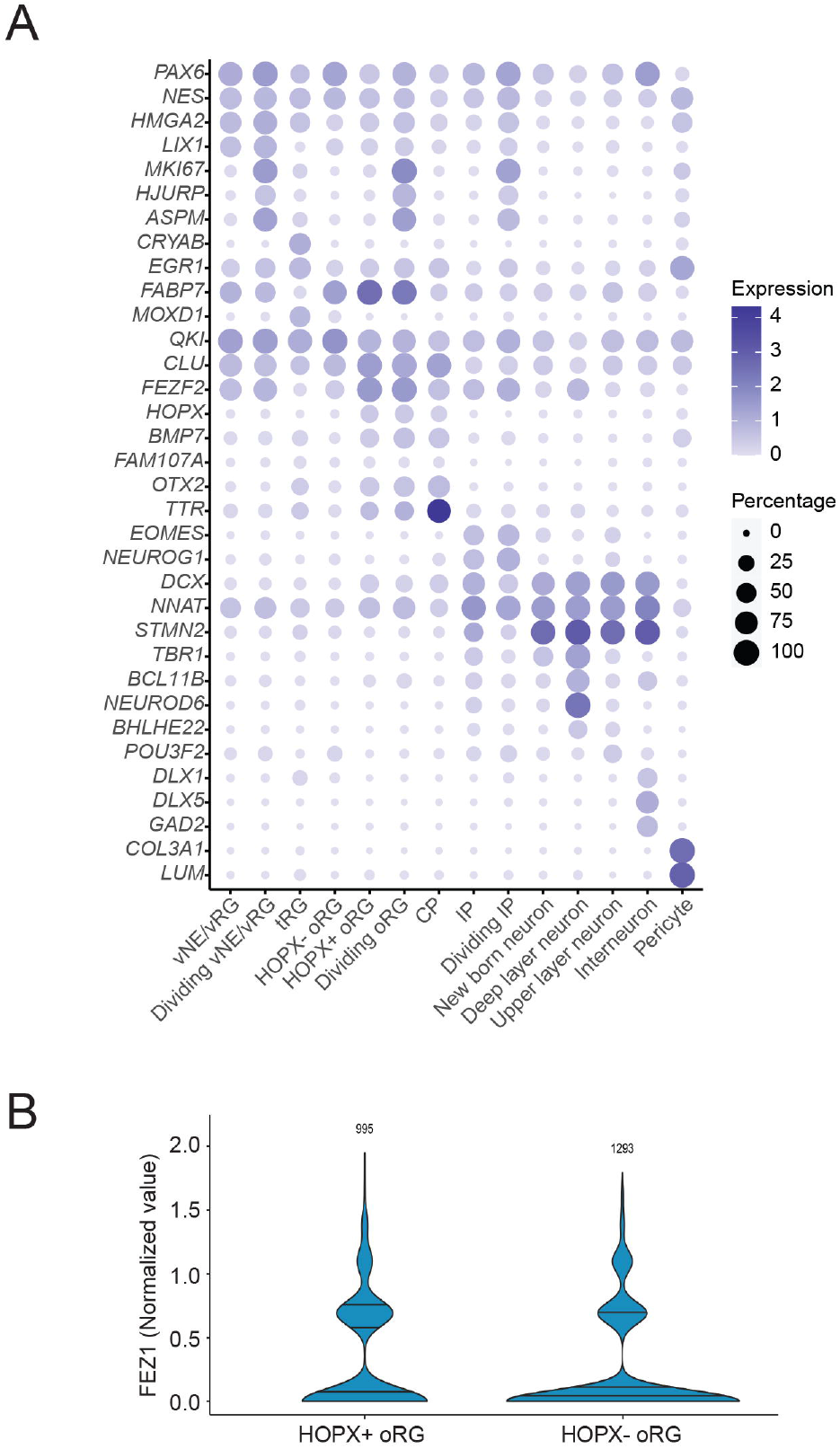
High throughput transcriptional analysis on D10 and D28 WT and FEZ1-null hCOs. (A) Spot plot of marker gene expressions in each cell clusters as identified by scRNA-seq on D28 WT and FEZ1-null hCOs and analyzed by Bioturing. Spot color scale indicated average log_2_ normalized value for each gene and spot size represented percentage of cells in each cluster expressed the corresponding gene. (B) Violin plot of cells expressing *FEZ1* in HOPX^+^ and HOPX^−^ oRG clusters. Cells were analyzed in pooled WT and FEZ1-null organoid samples. Quantiles were indicated in the plot, which was analyzed and generated by Bioturing.

**Figure S4.**
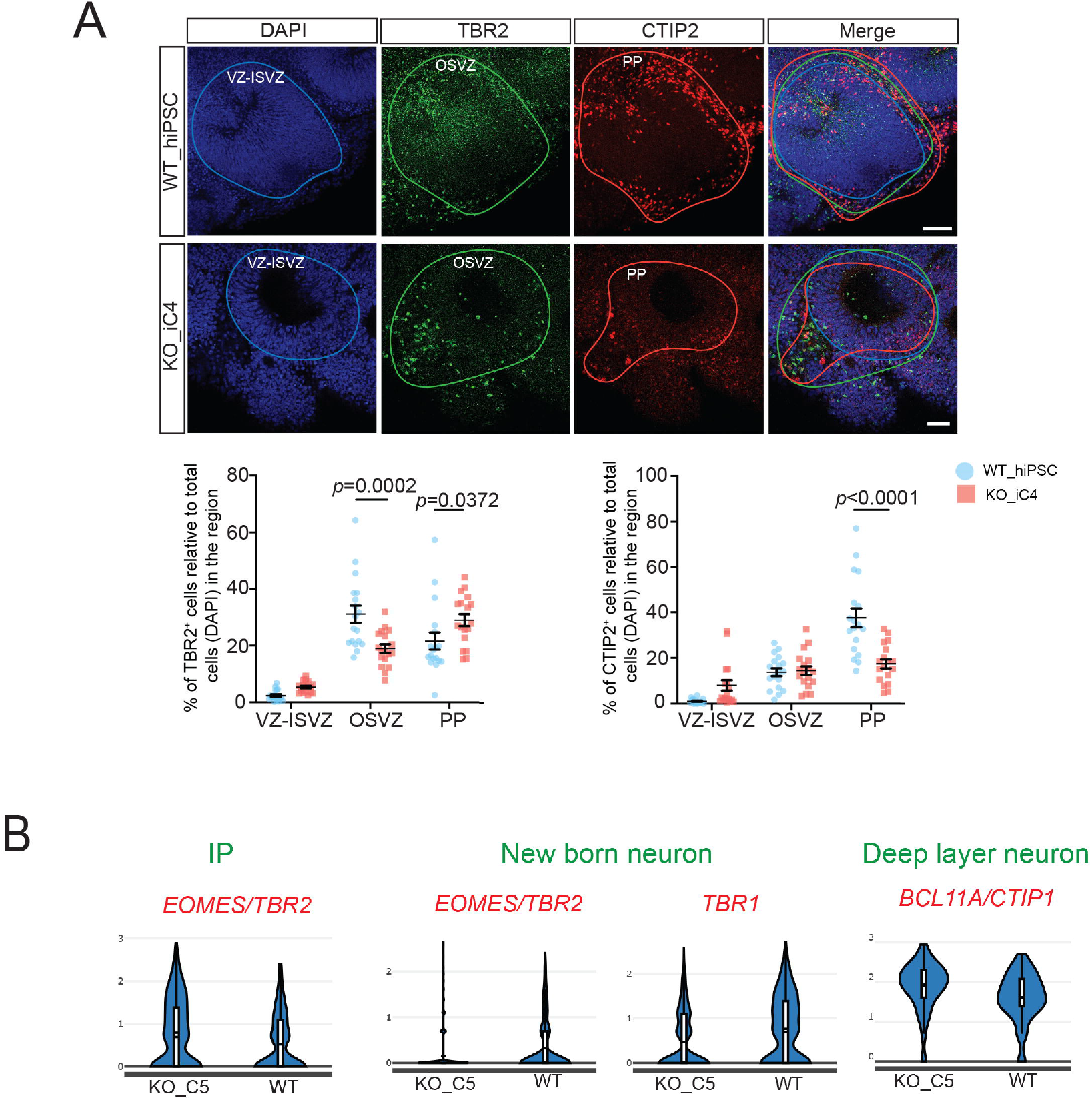
Lamination defects in FEZ1-null hCOs. (A) IF staining of TBR2 and CTIP2 showed that the abnormal lamination in D28 FEZ1-null hCOs was reproducible in the hiPSC_iC4 FEZ1-null clone. Layering in WT hCOs was unaffected. Scale bar: 50 μm. Lower panel shows quantification of TBR2^+^ and CTIP2^+^ cell populations in OSVZ and PP regions of D28 WT and FEZ1-null hiPSC hCOs. Values represent mean ± SEM (n= 3 independent organoid differentiations with total 6 organoids analyzed for WT and FEZ1-null groups, respectively. Each data point represents one analyzed organoid region, at least 3 regions were analyzed within each organoid, two-way ANOVA with Šidák’s multiple comparison is used to determine the indicated *p* values). (B) Violin plots of transcription factors (TFs) representative of IP, newborn neuron and deep layer neuron cell clusters with significant differential expression (analyzed by Bioturning) between WT and FEZ1-null hCOs.

**Figure S5.**
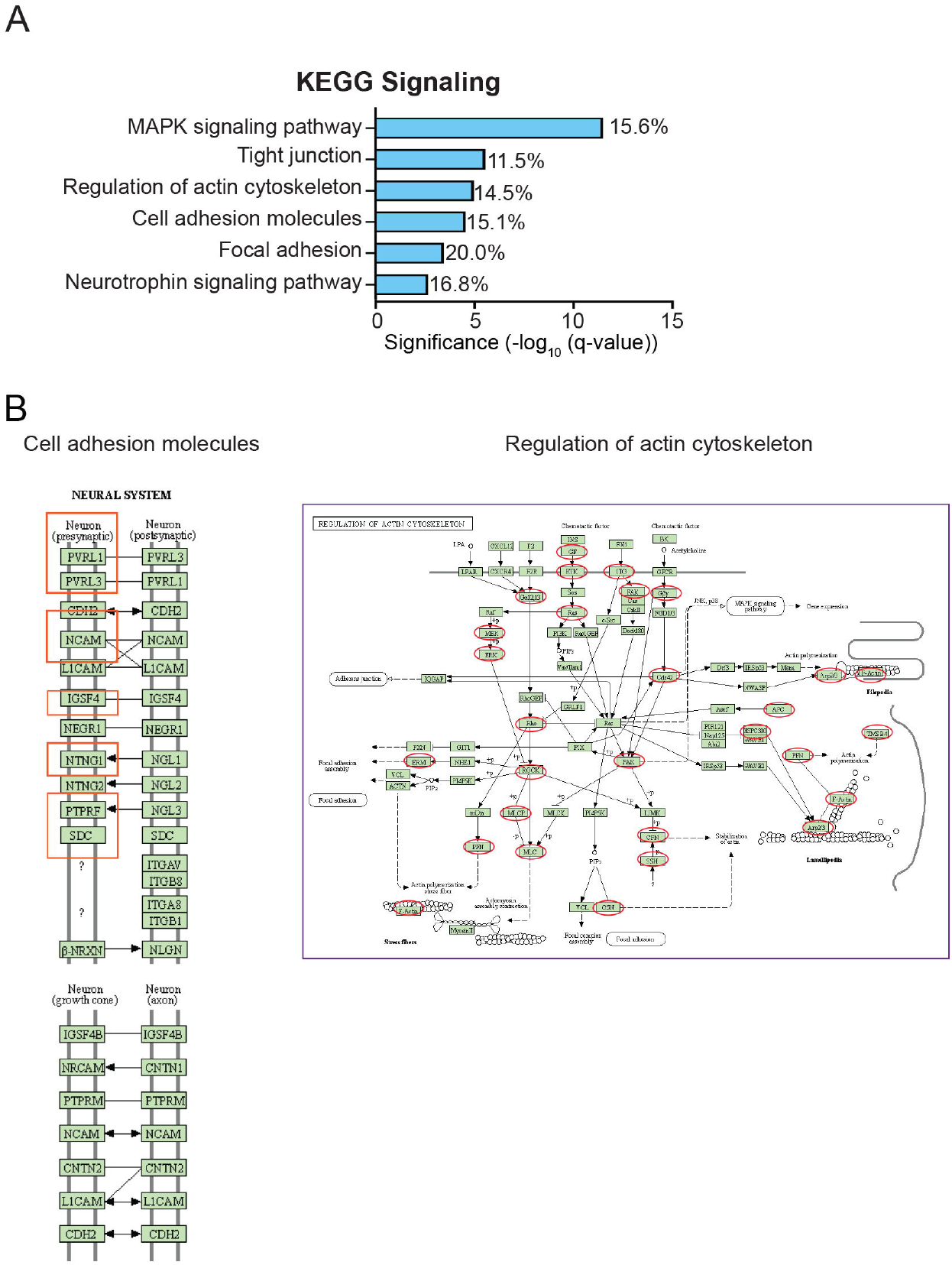
Impaired cell adhesion signaling and actin cytoskeleton regulation in D10 FEZ1-null hCOs. (A) KEGG signaling pathways involving DEGs between HOPX^+^ oRG and HOPX^−^ oRG as revealed by g:Profiler. (B) DEGs in HOPX^−^ oRG compared with HOPX^+^ oRG clusters revealed impairment on KEGG pathways (analyzed by DAVID) of cell adhesion molecules and regulation of actin cytoskeleton. Encircled red rectangles and ovals highlighted the DEGs presented in these pathways.

**Figure S6.**
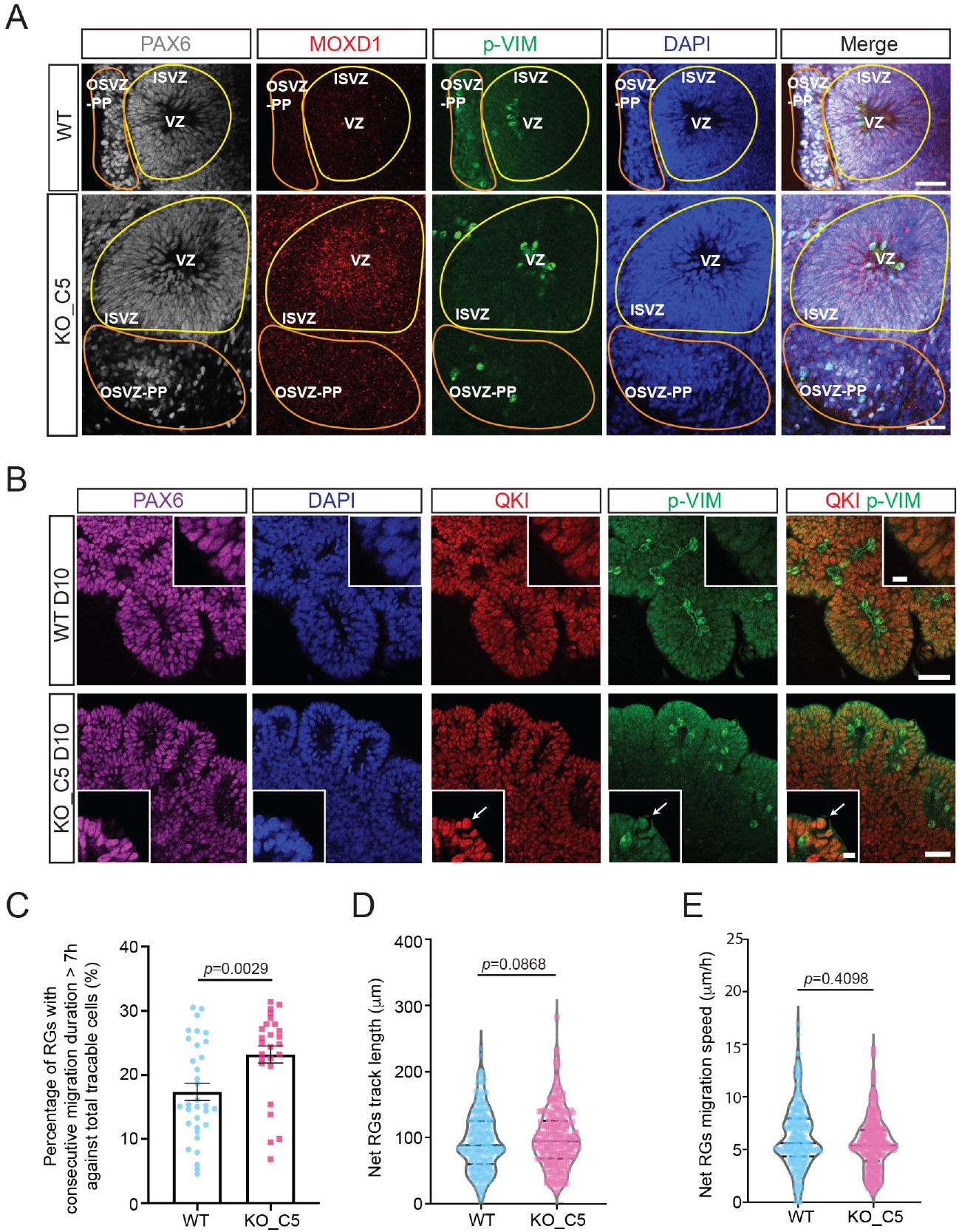
Ectopic localization of HOPX^−^ oRG and atypical cell migration in FEZ1-null hCOs. (A) IF staining of PAX6 (RG marker), MOXD1 (vRG and HOPX^−^ oRG marker) and p-VIM (mitotic RG marker) in D28 hCOs for both WT and FEZ1-null groups. More MOXD1^+^ oRG could be observed at OSVZ-PP in FEZ1-null hCOs. Scale bar: 50 μm. (B) IF staining of PAX6 (RG marker), QKI (vRG and HOPX^+^ oRG marker) and p-VIM (mitotic RG marker) in D10 hCOs for both WT and FEZ1-null groups. Upper right panel in WT D10 and lower left panel in KO D10 are the ROI images. Colocalization of QKI^+^ RG and p-VIM^+^ mitotic RG at outer surface of VZ/SVZ (white arrow) could be observed in D10 FEZ1-null hCOs. Scale bar: 50 μm and 10 μm (ROI). (C) Percentage of GFP^+^ RGs displayed consecutive migration behavior for more than 7 hours against total traceable cells in WT and FEZ1-null hCOs, respectively. Values represent mean ± SEM. (n=3 independent organoid differentiations, each dot represents value of one traced region, at least 8 regions were analyzed within each replicate, unpaired t-test with Welch’s correction with two-tailed *p* values indicated). Net RGs track length (D) and migration speed (E) in WT and FEZ1-null hCOs, respectively. Values represent as violin plot with median and quartiles indicated in dashed line. (n=3 independent organoid differentiations, each dot represents value of one traced cell, at least 100 cells were analyzed within each replicate, unpaired t-test with Welch’s correction with two-tailed *p* values indicated). VZ: ventricular zone; ISVZ: inner sub-ventricular zone; OSVZ-PP: outer sub-ventricular zone-preplate.

**Figure S7.**
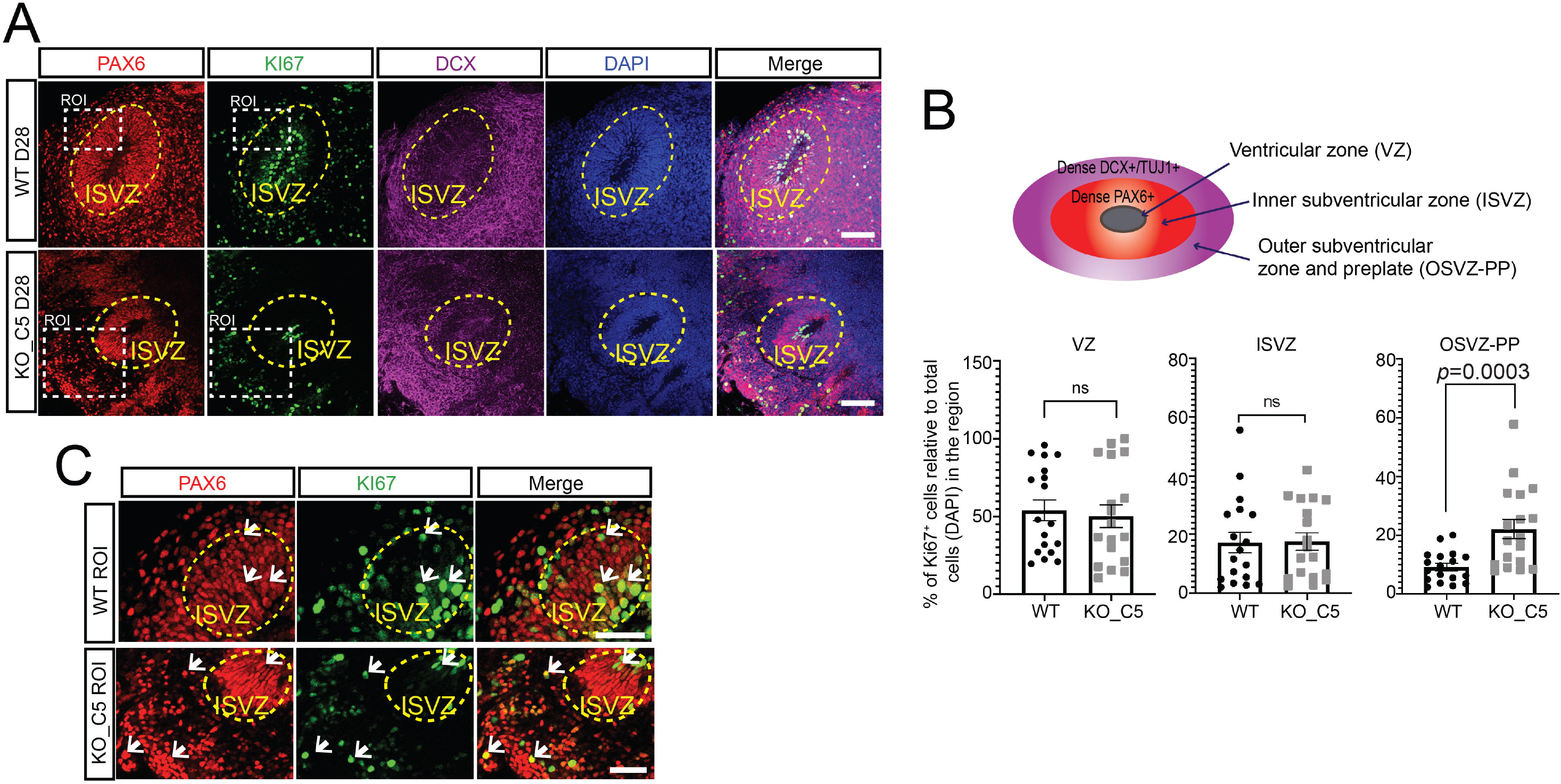
Ectopic localization and proliferation of RG in FEZ1-null hCOs. (A) IF staining of KI67, PAX6 and DCX showed mitotic cell localization in both D28 WT and FEZ1-null hCOs. Scale bar: 100 μm. (B) Quantification of KI67^+^ cells at VZ, ISVZ and OSVZ-PP in D28 hCO showed more mitotic cells located at OSVZ in FEZ1-null hCOs. Cartoon illustrates the identification of VZ-ISVZ and OSVZ-PP regions based on PAX6 and DCX IF staining. Values represent mean ± SEM. (n=3 independent organoid differentiations with total 6 organoids analyzed for WT and FEZ1-null group, respectively. Each data point represents value of one analyzed region, at least 5 regions were analyzed within each replicate, unpaired t-test with Welch’s correction with two-tailed *p* values indicated). (C) Zoomed ROI in (A) to demonstrate some KI67^+^ cells were co-localized with PAX6^+^ RGs (arrow), indicating that some mitotic cells were oRGs. Scale bar: 50 μm. VZ: ventricular zone; ISVZ: inner sub-ventricular zone; OSVZ: outer sub-ventricular zone; PP: preplate.

