## Supplementary material for "FEZ1 participates in human embryonic brain development by modulating neuronal progenitor subpopulation specification and migration": Materials and Methods

**KEY RESOURCES TABLE**

| REAGENT or RESOURCE | SOURCE | IDENTIFIER |
| --- | --- | --- |
| Antibodies | | |
| Rabbit anti-Neural Cell Adhesion Molecule (NCAM)(1:120) | Merck | Cat#AB5032, RRID:AB_2291692 |
| Rabbit anti-TUJ1 (ß3-Tubulin) (1:300) | Synaptic Systems | Cat#302302, RRID:AB_10637424 |
| Guinea Pig anti-TUJ1 (ß3-Tubulin) (1:300) | Synaptic Systems | Cat#302304, RRID:AB_10805138 |
| Rat anti-Ctip2 (1:500) | Abcam | Cat#ab18465, RRID:AB_2064130 |
| Rabbit anti-Fez1 (IF: 1:400; WB: 1:1000) | In house | clone361 |
| Rabbit anti-GAPDH (WB: 1:1000) | Abcam | Cat#ab9485, RRID:AB_307275 |
| Alexa Fluor™ 647 Phalloidin (1:500) (F-actin) | Thermo Fisher Scientific | Cat#A22287, RRID:AB_2620155 |
| Deoxyribonuclease I, Alexa Fluor™ 488 Conjugate (1:500) (G-actin) | Thermo Fisher Scientific | Cat#D12371 |
| Guinea pig anti-Doublecortin (DCX) (1:600) | Merck | Cat#AB2253, RRID:AB_1586992 |
| DAPI (4',6-Diamidino-2-Phenylindole, Dihydrochloride) | Thermo Fisher Scientific | Cat#D1306, RRID:AB_2629482 |
| Mouse anti-N cadherin (1:100) | Abcam | Cat#ab19348, RRID:AB_444868 |
| Rabbit anti-E-cadherin (IF: 1:50) | Santa Cruz | Cat#sc-7870, RRID:AB_2076666 |
| Rabbit anti-Nestin (1:200) | Abcam | Cat#ab92391, RRID:AB_10561437 |
| Sheep anti-Pax6 (1:50) | R&D Systems | Cat#AF8150, RRID:AB_2827378 |
| Mouse anti-Lin28a (6D1F9) (IF 1:1000) | Cell Signaling | Cat#5930S, RRID:AB_1903976 |
| Rabbit anti-Nanog (IF: 1:500) | Cell Signaling | Cat#3580S, RRID:AB_2150399 |
| Mouse anti-SSEA4 (IF: 1:500) | Abcam | Cat#ab16287, RRID:AB_778073 |
| Mouse anti-Phosphorylated Vimentin (1:250) | MBL | Cat#D076-3, RRID:AB_592963 |
| Mouse anti-SOX2 (1:300) | R&D Systems | Cat#MAB2018, RRID:AB_358009 |
| Rabbit anti-TBR2 (1:300) | Abcam | Cat#ab23345, RRID:AB_778267 |
| Mouse anti-Ki 67 (1:100) | BD Transduction Laboratories™ | Cat#610969, RRID:AB_398282 |
| Rabbit anti-QKI (IF: 1:100) | Abcam | Cat#ab126742, RRID:AB_11129508 |
| Rabbit MOXD1 (IF: 1:150) | Sigma-Aldrich | Cat#HPA035740, RRID:AB_10610141 |
| Donkey anti-Mouse IgG (H+L) Highly Cross-Adsorbed Secondary Antibody, Alexa Fluor Plus 488 (1:500) | Thermo Fisher Scientific | Cat#A32766, RRID:AB_2762823 |
| Donkey anti-Sheep IgG (H+L) Cross-Adsorbed Secondary Antibody, Alexa Fluor 555 (1:500) | Thermo Fisher Scientific | Cat#A-21436, RRID:AB_2535857 |
| Goat anti-Guinea Pig IgG (H+L) Highly Cross-Adsorbed Secondary Antibody, Alexa Fluor 555 (1:500) | Thermo Fisher Scientific | Cat#A-21435, RRID:AB_2535856 |
| Donkey anti-Rabbit IgG (H+L) Highly Cross-Adsorbed Secondary Antibody, Alexa Fluor Plus 647 (1:500) | Thermo Fisher Scientific | Cat#A32795, RRID:AB_2762835 |
| Goat anti-Mouse IgG (H + L)-HRP Conjugate (1:4000) | Bio-Rad | Cat#170-6516, RRID:AB_11125547 |
| Goat anti-Rabbit IgG (H + L)-HRP Conjugate (1:4000) | Bio-Rad | Cat#170-6515, RRID:AB_11125142 |
| Donkey anti-Rat IgG (H+L)(Cy2) (1:200) | Jackson ImmunoResearch | Cat#712-225-150, RRID: AB_2340673 |
| Donkey Anti-Guinea Pig IgG (H+L) (Cy2) (1:200) | Jackson ImmunoResearch | Cat#706-225-148, RRID: AB_2340467 |
| Donkey Anti-Sheep IgG (H+L) (Cy5) (1:200) | Jackson ImmunoResearch | Cat#713-175-147, RRID: AB_2340730 |
| Bacterial and Virus Strains | | |
| Lentiviruse (pLenti-CRISPR-FEZ1-KO ) | In house | N.A. |
| AAV (EF1a/CMV-GFP) | Gift from Dr.Ayumu Tashiro (NTU) | N.A. |
| Chemicals, Peptides, and Recombinant Proteins | | |
| Matrigel® hESC-Qualified Matrix, LDEV-free | Corning | Cat#354277 |
| TRIzol™ Reagent | Thermo Fisher Scientific | Cat#15596026 |
| Pierce™ 16% Formaldehyde (w/v), Methanol-free | Thermo Scientific | Cat#28908 |
| Matrigel® Growth Factor Reduced (GFR) Basement Membrane Matrix, LDEV-free | Corning | Cat#354230 |
| N-2 Supplement (100X) | Thermo Fisher Scientific | Cat#17502048 |
| B-27™ Supplement | Thermo Fisher Scientific | Cat#17504044 |
| Y-27632 | STEMCELL Technologies | Cat#72304 |
| Puromycin | Merck | Cat#P8833-10MG, CAS number: 58-58-2 |
| Critical Commercial Assays | | |
| CloneR™ | STEMCELL Technologies | Cat#05888 |
| CellTiter 96® AQueous One Solution Cell Proliferation Assay (MTS) | Promega | Cat#G3582 |
| CyQUANT™ XTT Cell Viability Assay | Thermo Fisher Scientific | Cat#X12223 |
| RapiClear 1.52 | SunJin Lab Co. | Cat#RC152001 |
| Taq PCR Core Kit | Qiagen | Cat#201223 |
| Wizard(R) Genomic DNA Purification Kit | Promega | Cat#A1120 |
| PowerUp™ SYBR™ Green Master Mix | Thermo Fisher Scientific | Cat#A25742 |
| STEMdiff™ Cerebral Organoid Kit | STEMCELL Technologies | Cat#08570 |
| Click-iT™ Plus EdU Alexa Fluor™ 555 Imaging Kit | Thermo Fisher Scientific | Cat#C10638 |
| RNeasy Mini Kit | Qiagen | Cat#74106 |
| SensiFAST^TM^ cDNA synthesis kit | Bioline | Cat#BIO-65054 |
| Experimental Models: Cell Lines | | |
| hESC H1 | WiCell | Cat#WA01 |
| AAVS-DCX-GFP IPSC | XCell Science | Cat#IP-001-ZIX4-1V |
| Oligonucleotides | | |
| Primers for RT-PCR: human Fez1: forward 5'-ACTACAACGCCAAGACC-3' and reverse 5'-AGAGCATCCCAAACCT-3'; human Gapdh: forward 5'-TGCACCACCAACTGCTTAGC-3' and reverse 5' GGCATGGACTGTGGTCATGAG-3' | In house | N.A. |
| Primers for PCR: human Fez1: forward 5'-ATAAACTCATCCTGAAAGTCGCTG-3' and reverse 5'-CCTCGTCCTGAAGGGTCTCCT-3' | In house | N.A. |
| Software and Algorithms | | |
| ImageJ/Fiji | (Schindelin et al., 2012) | <https://imagej.nih.gov/ij/download.html>  RRID:SCR_003070/ RRID:SCR_002285 |
| ZEN lite | Zeiss | <https://www.zeiss.com/microscopy/int/downloads.html>  RRID:SCR_013672 |
| Imaris 9.5.1 | Oxford Instruments | <http://www.bitplane.com/imaris/imaris>  RRID:SCR_007370 |
| BioTuring Browser | BioTuring | https://bioturing.com/bbrowser |
| Partek Flow | Partek | https://www.partek.com/partek-flow/ |

**DATA AND CODE AVAILABILITY**

Bioinformatics code are available upon reasonable request.

**EXPERIMENTAL MODEL AND SUBJECT DETAILS**

**Human ESC line and iPSC line (hPSC lines)**

All human stem cell work was performed with approval from National University of Singapore (NUS). hESC H1 hiPSC-DCX (AAVS-DCX-GFP) line was characterized and provided by Prof Zeng Xianmin in kind (Pei et al., 2015). The hESC line H1 (WiCell, Madison, WI) and hiPSC-DCX were maintained in *Mycoplasma* free conditions with routine *Mycoplasma* test every six months. Feeder free hPSC lines were maintained on hESC qualified Matrigel coated cell culture plate in mTeSR™1 (STEMCELL Technologies, Cat#85850), passaged with ReLeSR™ (STEMCELL Technologies, Cat# 05872) or Gentle Cell Dissociation Reagent (STEMCELL Technologies, Cat#100-0485) and frozen in CryoStor® CS10 (STEMCELL Technologies, Cat#07930) according to the manufacturer’s recommendations.

**METHOD DETAILS**

**Generation of FEZ1 knockout (KO) hPSC lines by CRISPR/CAS9**

Design of sgRNA, construction and production of lentiviruses have been previously described (Chua et al., 2021; Gunaseelan et al., 2021). Briefly, gRNA sequence targeting exon 2 of human *FEZ1* gene (forward 5’-AATCAGCTTCAAGTCCATGG-3’ and reverse 5’-CCATGGACTTGAAGCTGATT-3’) were designed using the online CRISPR design tool (<http://crispr.mit.edu/>) and inserted into LentiCRISPRv2 plasmid (Addgene, plasmid #52961). pLenti-CRISPR-FEZ1-KO vector was co-transfected with pMDLg/pRRE (Addgene; plasmid #12251), pRSV-rev (Addgene; plasmid #12253) and pMD2.G helper plasmids (Addgene; plasmid #12259) (2:1:1) into HEK293. Cell supernatant containing lentiviruses was collected after 24 h of post-transfection and concentrated with Amicon Ultra-15 filters (Millipore) by ultracentrifugation at 3,220× *g* for 30 min at 4 ºC.

To generate FEZ1 KO hPSCs, cells at 60% to 70% confluence one day after seeding on 12-well plates were infected with 100 μL of concentrated lentiviruses and cultured in mTeSR™1 for 24 hours. Twenty-four hours post infection, successfully transduced cells were selected using 5 μg/ml (H1) or 2 μg/ml (iPSC-DCX) puromycin for 24 to 48 h. Positively selected cells were then dissociated with Accutase (ThermoFisher Scientific, Cat#A1110501) and passed through 40 μM cell strainer (Fisher scientific, Falcon, Cat#352340). Single cells obtained were plated at a density of 1,500 cells on a 10-cm dish coated with hESC qualified Matrigel. Cells were maintained in mTeSR™1 supplemented with CloneR™ (STEMCELL Technologies) for the first 3 days of culture followed by mTeSR™1 maintenance culture for another 5 to 8 days until individual colonies reached a diameter larger than 1 mm. After a 3-minute treatment with Dispase (STEMCELL Technologies, Cat#07923), colonies were individually picked up under an inverted microscope and carefully transferred into individual wells of 24-well plate. The selected clones were then expanded and stored to generate clonal banks of FEZ1 KO hPSCs for further characterization and application. Karyotyping for selected FEZ1 KO hPSCs was performed by the KK Women’s and Children’s Hospital (Singapore). Cell proliferation was measured using the CyQuant kit (ThermoFisher Scientific, Cat#X12223) according to the manufacturer’s recommendations.

**Genomic sequencing of** **FEZ1 knockout (KO) hPSC lines**

To confirm the successful generation of FEZ1 KO hPSC lines, selected clones from FEZ1-null hPSCs bank were expanded and cell DNA were extracted and purified with Wizard® Genomic DNA Purification Kit (Promega, Cat#A1120). DNA amplification prior to sequence was performed by Taq PCR Core Kit (Qiagen, Cat#201223) and FEZ1 primers targeting the editing site (forward 5'-ATAAACTCATCCTGAAAGTCGCTG-3' and reverse 5'-CCTCGTCCTGAAGGGTCTCCT-3'). Amplicons were analyzed by sequencing (forward 5’-AATCAGCTTCAAGTCCATGG-3’ and reverse 5’-CCATGGACTTGAAGCTGATT-3’) and results were analyzed with FinchTV and BLAST® (https://blast.ncbi.nlm.nih.gov/Blast.cgi) to characterize the nature of the indels in hPSCs.

**2D neural epithelium (NE) induction**

To differentiate NE on monolayer culture, hPSCs were dissociated with Accutase (ThermoFisher Scientific, Cat#A1110501) and plated on hESC qualified Matrigel (Corning, Cat#354277)-coated 6-well cell culture plates at a density of 0.1 million cells/cm^2^. Cells were maintained in freshly prepared N2B27 medium (50 ml of Neurobasal medium (ThermoFisher Scientific, Cat#21103049), 50 ml of DMEM/F12 Glutmax (ThermoFisher Scientific, Cat#10565018), 0.1 mM of β-mercaptoethanol (ThermoFisher Scientific, Cat#21985023), 0.2 mM of glutamine (ThermoFisher Scientific, Cat#25030081), 0.5× N2 supplement (ThermoFisher Scientific, Cat#17502048) and 0.5× B27 supplement (ThermoFisher Scientific, Cat#17504044) to obtain 100 ml N2B27 medium) with daily change of medium for 6 days as previously described (Ying et al., 2003). After 6 days of NE induction, the cells were harvested based on the requirements for various downstream assays. For each assay, at least three independent experiments were performed.

**Generation of human cerebral organoids (hCOs)**

hCOs were generated by using the STEMdiff™ Cerebral Organoid Kit (STEMCELL Technologies, Cat#08570), which consisted of Embryoid body (EB) formation media, induction medium, expansion medium and organoid maturation medium. Freshly thawed hPSC were maintained in mTeSR™1 until 70-80% confluency and then dissociated with Accutase to obtain single cells. EB were generated by seeding 9,000 viable cells/well in a 96-well round bottom ultra-low attachment microplate (Corning, Cat. 7007) in EB formation media supplemented with ROCKi/Y-27632 (STEMCELL Technologies, Cat#72304). After 5 days, EBs were transferred to individual wells containing induction medium in 48-well suspension culture plates (Greiner, Cat. 677102) and grown for 2 days. At day 7, each EB was embedded in Matrigel (Corning, growth factor reduced basement membrane matrix, Cat#354230) and transferred into a 6-well suspension culture plate (Greiner, Cat#657185) containing expansion medium. At day 10, expansion medium was removed from hCOs and replaced with organoid maturation medium. The culture plates containing the hCOs were then placed onto an orbital shaker (Stuart, Cat# SSM1) in a 37ºC incubator with gentle agitation at 80 rpm to allow hCOs to develop in organoid maturation medium with medium change every 3 or 4 days. Organoid morphologies were routinely monitored by microscopy (Nikon, Eclipse TS100 equipped DS-Fi1/Digital sight DS-L2 camera and Nikon, Eclipse TE2000-E). For each downstream assay, at least 3 or 4 biological repeats of organoids were used.

**Immunofluorescence (IF) microscopy**

For monolayer cell cultures, cells were fixed with 4% formaldehyde (by diluting the 16% formaldehyde (ThermoFisher Scientific, Cat#28908) with 1xPBS) for 20 mins at room temperature and then washed with calcium free phosphate-buffered saline (1×PBS, 1^st^ BASE, Cat#BUF-2040-1X500ml, containing 137 mM sodium chloride, 2.7 mM potassium Chloride and 12 mM phosphate buffer). Permeabilization was performed using 0.4% Triton X-100 (Sigma-Aldrich, Cat#X100-500ML) in 1×PBS for 20 minutes. After washing, cells were blocked in blocking buffer (2% BSA, 0.2% Triton X-100 in 1×PBS) for 1 hour. Incubation with primary antibodies were performed overnight at 4ºC. After washing, samples were incubated with secondary antibodies at room temperature for 2 hours. After washing, samples were mounted with FluorSave^TM^ (Merck, Cat#345789) for epifluorescence (Nikon, Eclipse Ti2; Zeiss, Axio Observer Z1 and Nikon, Eclipse TE2000-E) or confocal (Zeiss LSM800) imaging. For each assay, at least 3 independent experiments were performed.

Whole hCOs were fixed with 4% FA for 2 hours (D10 and D28 hBOs) or 3 hours (hCOs older than 28 days) at room temperature. Organoids were then washed with 1× PBS to remove any remaining FA. Fixed hCOs were either stored in 1× PBS at 4ºC or be permeabilized for 2 days (1% Triton X-100 and 1% DMSO in 1×PBS) and blocked for an additional 2 days (2% BSA, 1% Triton X-100, 1% DMSO and 1% sodium azide in 1× PBS). After blocking, organoids were then incubated with primary antibodies diluted in antibody dilution buffer (2% BSA, 0.2 % Triton X-100, 1% DMSO, 1% sodium azide and 1% Heparin in 1× PBS) for 2 to 3 days depending on the age of the hCOs. Samples were than washed thrice with washing buffer (3% NaCl and 0.2% Triton X-100 in 1× PBS) for at least 12 hours. hCOs were then incubated with secondary antibodies and DAPI in antibody dilution buffer for 2 to 3 days and washed as before. After the final washing step, hCOs were rinsed once with 1× PBS and left in 1× PBS for at least 6 hours to remove any remaining Triton X-100. hCOs were then cleared overnight with RapiClear 1.52 (Sunjin Lab, Cat#RC152001) and mounted on iSpacer (Sunjin Lab, 0.5 mm, Cat#IS008). Scanning of cleared hBOs was performed on Zeiss LSM800 confocal microscopy equipped with long working distance (0.57 mm) multi-immersion objective lens (Zeiss, Objective LD LCI Plan-Apochromat 25x/0.8 Imm Corr DIC M27). For each assay, at least 3 independent experiments were performed.

Antibodies used for IF labeling are as follows: Primary antibodies: Sheep anti-Pax6 (1:50, R&D Systems, Cat#AF8150); Mouse anti-N cadherin (1:100, Abcam, Cat#ab19348); Mouse anti-Phosphorylated Vimentin (1:250, MBL, Cat#D076-3); Rabbit anti-Nestin (1:200, Abcam, Cat#ab92391); Rabbit anti-TUJ1 (1:300, Synaptic Systems, Cat#302302); Guinea Pig anti-TUJ1 (1:300, Synaptic Systems, Cat#302304); Rat anti-Ctip2 (1:500, Abcam, Cat#ab18465); Guinea pig anti-DCX (1:600, Merck, Cat#AB2253); Rabbit anti-TBR2 (1:300, Abcam, Cat#ab23344); Mouse anti-Ki 67 (1:100, BD Transduction Laboratories™, Cat#610969); Rabbit anti-NCAM (1:120, Merck, Cat#AB5032); Rabbit anti-QKI (1:100, Abcam, Cat#ab126742). Secondary antibodies: Donkey anti-Mouse IgG (H+L) Highly Cross-Adsorbed Secondary Antibody, Alexa Fluor Plus 488 (1:500, Thermo Fisher Scientific, Cat#A32766); Donkey anti-Sheep IgG (H+L) Cross-Adsorbed Secondary Antibody, Alexa Fluor 555 (1:500, Thermo Fisher Scientific, Cat#A-21436); Donkey anti-Rabbit IgG (H+L) Highly Cross-Adsorbed Secondary Antibody, Alexa Fluor Plus 647 (1:500, Thermo Fisher Scientific, Cat#A32795); Goat anti-Guinea Pig IgG (H+L) Highly Cross-Adsorbed Secondary Antibody, Alexa Fluor 555 (1:500, Thermo Fisher Scientific, Cat#A-21435); Donkey anti-Rat IgG (H+L)(Cy2) (1:200, Jackson ImmunoResearch, Cat#712-225-150).

**EdU labeling**

EdU assay was performed using the Click-iT^®^ Plus EdU Alexa Fluor™ 555 Imaging Kit (ThermoFisher Scientific, Cat#C10638). Day 23 hCOs were first incubated with 5 μM EdU in maturation medium with continuous agitation. After 4 hours of incubation, EdU medium was removed and replaced with fresh maturation medium after rinsing twice with DMEM/F-12 (ThermoFisher Scientific, Cat#11330107). Edu-labelled hCOs were then fixed at day 28 with 4% FA for 2 h at room temperature. Immediately after fixation, permeabilization and blocking were performed as described in the preceding section. After blocking, EdU-labelled hCOs were incubated overnight at room temperature protected from light with Click-iT® Plus reaction cocktail containing 1X Click-iT® reaction buffer, copper protectant, Alexa Fluor® picolyl azide and reaction buffer additive. After washing, hCOs were incubated again in blocking buffer overnight followed by primary and secondary antibody staining steps (details can be found under section on “Immunofluorescence (IF) microscopy”) for hCOs to co-stain EdU with other antibodies. Antibodies used for co-staining with BrdU were as follows: Primary antibodies: Sheep anti-Pax6 (1:50, R&D Systems, Cat#AF8150); Guinea Pig anti-TUJ1 (1:300, Synaptic Systems, Cat#302304). Secondary antibodies: Donkey Anti-Guinea Pig IgG (H+L) (Cy2) (1:200, Jackson ImmunoResearch, Cat#706-225-148) and Donkey Anti-Sheep IgG (H+L) (Cy5) (1:200, Jackson ImmunoResearch, Cat#713-175-147). Four independent experiments were performed and analyzed for D28 BrdU assays.

**Protein sample preparation and immunoblot assays**

For monolayer cell cultures, cells were lysed with HEPES lysis buffer (50 mM HEPES, 150 mM NaCl, 1 mM EDTA, 1% Triton X-100, pH 7.2) as described previously (Butkevich et al., 2016). In brief, cells were washed once with cold 1xPBS and subsequently incubated with cold HEPES buffer on rocker for 10 minutes at 4°C. Cell lysates were collected and centrifuged at 10,000× *g* for 10 minutes at 4°C. Supernatant was collected and mixed with 4×NuPAGE LDS Sample Buffer (ThermoFisher Scientific, Cat#NP0008) for immunoblot analyses or stored at -20ºC.

For organoid samples, hCOs were homogenized in sucrose buffer (320 mM sucrose, 1 mM EDTA, 5 mM HEPES, pH 7.4) with homogenizer (Bandelin, Sonopuls ultrasonic homogenizer) to obtain the homogenized cell lysate. Homogenates were centrifuged at 2,000× *g* for 2 minutes to separate nuclei (pellet) from cell lysate (Pavlos et al., 2010). The S1 supernatants were then centrifuged again at 14,500× *g* for 12 minutes. The S2 supernatant were then harvested and mixed with 4×NuPAGE LDS Sample Buffer for immunoblot analyses or stored at -80ºC.

To perform immunoblot analysis, protein samples were resolved by SDS-PAGE and transferred onto nitrocellulose membranes (Bio-Rad) using the Trans-Blot Turbo Transfer System (Bio-Rad). Membranes were then blocked with 5% skim milk in TBST (1.5 M NaCl, 0.5% Tween 20, 150 mM Tris-HCl, pH 7.4) and subsequently incubated with primary antibodies at 4 ºC overnight. After three washes with TBST, the membranes were then incubated with secondary antibodies for 1 hour at room temperature. After washing, membranes were treated with Immobilon Forte Western HRP Substrate (Millipore, Cat#WBLUF0500). Protein bands were visualized and captured by Azure Biosystem C300 (Azure Biosystems). Quantification of immunoblot were performed with ImageJ/Fiji and the intensities normalized against GAPDH. At least 3 independent batches of protein samples were used to run the immunoblot assay and quantification.

Antibodies used for immunoblot assays are as follows: Primary antibodies: Rabbit anti-Fez1 (IF: 1:1000); Rabbit anti-GAPDH (WB: 1:1000, Abcam, Cat#ab9485). Secondary antibodies: Goat anti-Rabbit IgG (H + L)-HRP Conjugate (1:4000, Bio-Rad, Cat#170-6515).

**Cell migration studies in hCOs**

To observe cell migration in unfixed hCOs, 5 day-old hCOs were transferred to cell culture imaging dishes (ibidi, Cat#81156) in induction medium. At day 6, hCOs were infected with GFP-expressing AAVs (a kind gift from Dr Ayumu Tashiro) for 24 hours in induction medium. Live imaging was carried out by confocal microscopy on a Zeiss LSM800 with the temperature of the imaging chamber set at 37ºC. For imaging, organoid maturation medium was supplemented with 25 mM HEPES to maintain pH. Images for the GFP and differential interference contrast channels were acquired at a rate of 1 frame every 20 minutes over a 20-hour recording period. At least 10 different locations were chosen for analyses per hCO. 3 independent experiments were performed for each group of hCOs for live imaging. Image stacks were processed and analyzed using Imaris 9.5.1(Bitplane). The details of quantification are described in the section on “Quantification of IF images”.

**Bulk RNA sequence (RNA-seq) sample preparation and RT-PCR**

Cells or hCOs were lysed with the TRIzol™ reagent (ThermoFisher Scientific, Cat#15596026). The lysates were then mixed with chloroform and centrifuged at 16,200× *g* for 15 minutes at 4 ºC. The top colorless layer was carefully aspirated, mixed with 70% ethanol and loaded into RNeasy spin columns (Qiagen, RNeasy Mini Kit, Cat#74106) to obtain purified mRNA.

For bulk RNA sequence, a pre-quality check was done by agarose electrophoresis to check the quality of the 18S and 28S ribosomal RNAs. Samples were then processed by Novagene (Singapore) for additional QC, library preparation and total RNA-seq using an Illumina NovaSeq 6000 platform to obtain a sequencing depth of 20 million read pairs per sample. Data were acquired from 3 independent batches of samples with total 4 repeats for both WT and FEZ1-null groups.

For RT-qPCR assays, 1 μg of mRNA was converted to cDNA using the SensiFAST^TM^ cDNA synthesis kit (Bioline, Cat#BIO-65054). 10 ng of cDNA were used for PCR amplification of each targeted gene using the PowerUp™ SYBR™ Green Master Mix (ThermoFisher Scientific, Cat#A25742). RT-qPCR quantification was performed on QuantStudio 3 Real-Time PCR System (ThermoFisher Scientific). Primers used in RT-PCR study were as follows: human *Fez1*: forward 5'-ACTACAACGCCAAGACC-3' and reverse 5'-AGAGCATCCCAAACCT-3'; human *Gapdh*: forward 5'-TGCACCACCAACTGCTTAGC-3' and reverse 5' GGCATGGACTGTGGTCATGAG-3'. The expression levels of genes were normalized against *Gapdh*. Transcript levels from differentiated samples were compared against undifferentiated cells to obtain fold change. Repeats were conducted with 3 independent batches of samples.

**Single cell RNA sequencing (scRNA-seq)**

We modified our in-house rat hippocampal cell isolation protocol to obtain single cell samples for scRNA-seq (Chua et al., 2021). hCOs were dissected into smaller pieces by scalpel and trypsinized by Trypsin/EDTA solution (Lonza, Cat#CC-5012) for 25 min. Digestion was stopped by adding serum medium (Modified Eagle’s Medium (Sigma-Aldrich, Cat#M2414-500ML) supplemented with 5% fetal calf serum (Biochrom, Cat#50115), 2 mM L-alanyl-L-glutamine (Biochrom, Cat#K0302), 1× MEM Vitamin Solution (Biochrom, Cat#K0373), 0.2x Mito+Serum extender (BD, Cat#355006) and 21 mM D-glucose (Merck, Cat#1.08337.1000). The fragments were then gently triturated with a Pasteur pipette and the suspension spun down at 500× *g* for 5 min. Pelleted cells were resuspended in ESCAPE^TM^ buffer (provided by Proteona) and passed through a 40 μm strainer. Cell viability and counts were determined (viability > 85% and cell count > 1 million/sample). Single cell sample library preparation and sequencing were done by Proteona (Singapore). Only QC passed single alive cells were used for library preparation using the 3’RNA V3 kit on 10× Genomics Chromium Controller. Cell suspensions were processed with the Chromium Controller (10x Genomics) using Chromium Single Cell 3’ Gem, Library & Gel Bead Kit v3 (10x Genomics, Cat# PN-1000078) and Chromium Single Cell B Chip Kit (10x Genomics, Cat# PN-1000074). RNA libraries that met QC criteria were sent for further sequencing using HiSeq X (Illumina) with 150 bp paired end reads.

**Bulk RNA-seq analysis**

The raw RNA-seq data was uploaded to NUS Cancer Science Institute of Singapore (CSI) NGS Portal (<https://csibioinfo.nus.edu.sg/csingsportal>) (An et al., 2020) to perform gene expression profiling and differential expression analysis. Briefly, clean reads were aligned to the reference human genome (hg19) with default parameters (Dobin et al., 2013). The gene expression quantification was done by using HTSeq-count in strand-specific mode with “-s reverse” option and read counts only from the sense strand were used for each gene (Anders et al., 2015). Differential gene expression analysis was performed by using DESeq2 (Love et al., 2014). The genes that did not express or in low expression (read counts less than 2 on average per sample) were removed from the analysis. After correction for multiple hypothesis testing (adjusted p value (Padj) < 0.05, using Benjamini-Hochberg method), 87 and 439 genes were identified with significant up- and down-regulation in the FEZ1-null group when compared to WT group. The differentially expressed genes (DEGs) were performed Gene Ontology (GO) and pathway enrichment analysis by Metascape (Zhou et al., 2019) and disease related gene enrichment analysis was run on Enrichr (<https://maayanlab.cloud/Enrichr/>).

**scRNA-seq analysis**

The Cell Ranger 3.1.0 pipeline (10x Genomics) was used to align reads from scRNA-seq to the GRCh38 human reference genome and the associated gene by cell count matrix was generated. The default parameters were used except for specifying the expected number of cells (‘--expect-cells=10000’). The resulting matrices containing the unique molecular identifier (UMI) counts were merged from WT and FEZ1-null samples into a single object and analyzed by the Seurat R package v.3.1 (Butler et al., 2018). Cells expressing a range of 1000-8000 genes and less than 20% of mitochondrial genes were retained based on the inspection of feature distributions, and genes expressed in less than 10 cells were filtered out. UMI counts were normalized and variance-stabilized by using ‘SCTransform’ function which employs a regularized negative binomial regression to remove technical differences between cells due to sequencing depth (Hafemeister and Satija, 2019). This step was accompanied by regressing out variation due to proportion of mitochondrial gene expression and identifying top 3000 most variable genes. Principal component analysis (PCA) was performed on the normalized data for the variable genes, and the first 30 principal components were used for clustering. In this PCA space, the cells were clustered by finding the nearest 20 neighbors to each cell (FindNeighbors function with ‘dims=1:30’), building a Shared Nearest Neighbor (SNN) graph with edges between neighbor cells weighted by the Jaccard index, and performing Louvain algorithm on the resulting graph (FindClusters function with ‘resolution=0.8’). The clusters of cells were then visualized in two-dimensional space by using t-distributed Stochastic Neighbor Embedding (t-SNE) projection. The resulting 26 clusters were manually merged into 14 clusters and assigned to corresponding cell types based on the expression of known marker genes: vNE/vRG (ventricular neuroepithelium/ radial glia cells ): *LIX1, NES, HMGA2, PAX6*; Dividing vNE/vRG: *MKI67, LIX1, PAX6, ASPM*; tRG (truncated radial glia cells): *CRYAB, EGR1, HMGA2*; HOPX- oRG (HOPX- outer radial glia cells): *FABP7, MOXD1, QKI, CLU*; HOPX+ oRG (HOPX+ outer radial glia cells): *FABP7, CLU, FEZF2, HOPX*; Dividing oRG: *MIK67, HJURP, FABP7, CLU, FEZF2, HOPX*; CP (choroid plexus): *TTR, OTX2*; IP (intermediate progenitor): *EOMES, DCX, BEUROG1*; Dividing IP: *EOMES, MKI67, BEUROG1*; New born neuron: *STMN2, DCX*; Deep layer neuron: *STMN2, NEUROD6, BCL11B*; Upper layer neuron: *POU3F2, BHLHE22*; Interneuron: *DLX5, GAD2, DLX1*; Pericyte: *COL3A1, LUM*.

For each sample (WT and FEZ1-null), differentially expressed genes (DEGs) that were upregulated or downregulated in each cluster compared to the rest of the cells were identified and visualized by using Partek Flow (https://www.partek.com/partek-flow/) and Bioturing (<https://bioturing.com/>) (DEG filtering: false discovery rate (FDR)<0.01, log_2_FC>0.137 or <-0.137)(Le et al., 2020). Gene Ontology (GO) and pathway enrichment analysis of DEG in specific clusters were visualized by Metascape (Zhou et al., 2019), DAVID and g:Profiler (version e99_eg46_p14_f929183) with g:SCS multiple testing correction method, applying a significance threshold of 0.05(Raudvere et al., 2019).

**F-actin and G-actin assays**

F-actin and G-actin measurement was performed as described (Huang et al., 2020). D28 hCOs were dissociated into single cells using the same method used for scRNA-seq. Harvested cells were resuspended in organoid maturation medium and passed through a 40 μm filter strainer to obtain single cells. Cells then seeded at a density of 0.05 million/well on 24-well size coverslips coated with hESC qualified Matrigel. After two days in culture, cells were fixed with 4% FA for 15 min. Permeabilization and blocking of the samples are performed as mentioned in the section on immunofluorescence microscopy. After blocking, cells were stained with antibody Alexa Fluor™ 647 Phalloidin (F-actin, ThermoFisher Scientific, Cat#A22287), Deoxyribonuclease I, Alexa Fluor™ 488 Conjugate (G-actin, ThermoFisher Scientific, Cat# D12371) and DAPI (ThermoFisher Scientific, Cat#D1306) for 2h at room temperature. Cells were then washed thrice in 1×PBS and mounted with FluorSave^TM^ (Merck, Cat#345789) for imaging (Nikon, Eclipse Ti2; Zeiss, Axio Observer Z1 and Nikon, Eclipse TE2000-E). The same exposure settings were used for acquiring images from WT and FEZ1-null group. Images were analyzed by with ImageJ/Fiji with quantification details in the following section. At least three independent experiments were performed.

**Quantification of IF images**

For quantification of F-actin and G-actin (Figure 4E), ImageJ/Fiji was used to obtain fluorescent intensities using the raw integrated density (RawIntDen) measurement under Analyze for both F- and G-actin channels. The G- to F-actin ratios for WT or FEZ1-null hCOs were normalized to the mean WT G- to F-actin ratio for each batch. At least 10 images were acquired from independent fields per group per batch. Data from 3 independent experiments were used to generate the plot.

For imaging of whole hCOs, 3D image analyses were performed using Imaris 9.5.1. For NCAM quantification (Figure S4D), the filament function was used to trace intact NCAM positive processes to obtain neurite length. A total of 8 hCOs from 4 independent experiments were analyzed for each genotype.

For quantification of p-VIM and QKI (Figure 6A and B), the spots function in Imaris 9.5.1 was used with estimated diameter of nuclei at 6 μm and intensity mean channel of p-VIM^+^ or QKI^+^ to filter and generate p-VIM^+^ or QKI^+^ spots, which could reflect the corresponding p-VIM^+^ or QKI^+^ cells for counting. Total cell numbers were obtained by counting DAPI^+^ cells using spots function. The localization and number of p-VIM^+^ or QKI^+^ cells were grouped as either adjacent to the ventricular zone (VZ) or in subventricular zone (SVZ). The ratios of p-VIM^+^ or QKI^+^ cells relative to total cell numbers were calculated by number of p-VIM^+^ or QKI^+^ cell divided by total cell number in each image. At least 5 different regions were analyzed within one organoid. A total of 8 hCOs from 3 independent experiments were used to perform the quantification for each genotype.

For analyses of Edu, Ki67 and the neuronal layers (Figure7 and S7), rosette layers were defined as follows (Kostovic, 2020; Molnar et al., 2019; Pollen et al., 2015; Qian et al., 2016). Dense PAX6^+^ region was identified as VZ-ISVZ while the DCX^+^ or TUJ1^+^ region that was distant from VZ was labelled as OSVZ-PP. TBR2^+^ cell region in WT was chosen as OSVZ while the region widths were used to define the OSVZ area in FEZ1-null genotype. The region outside OSVZ, where cells showed CTIP2^+^ was determined as PP. Accordingly, each region was manually drawn and the spots function in Imaris 9.5.1 was used to calculate the number of corresponding cells for the respective cell marker in each region and to normalized against total number (DAPI^+^) of cells (with estimated diameter of nuclei at 4 μm and intensity mean channel of DAPI^+^, PAX6^+^, TBR2^+^, CTIP2^+^, EdU^+^ and Ki67^+^ to filter, generate and calculate corresponding spots). At least 5 regions were analyzed within each organoid. For layering analyses, a total of 9 hCOs from 4 independent experiments (hESC H1 hCOs) and a total of 6 hCOs from 3 independent experiments (hiPSC hCOs) per genotype were analysed. For EdU and Ki67 quantifications, a total of 6 hCOs were analysed per genotype. VZ: ventricular zone; ISVZ: inner sub-ventricular zone; OSVZ: outer sub-ventricular zone; PP: preplate.

**Quantification of cell migration**

Migration of GFP+ cells recorded by live imaging was analyzed by the tracking surface over time function in Imaris 9.5.1. In brief, objective with detail of 1.25 μm was used to help identify GFP^+^ cells. Autoregressive motion with maximum gap size of 3 μm and a distance of 10 μm were used for tracking. Based on these settings, total traceable cell counts were collected in each location. Only cells with more than 7 hours of consecutive migration were used for further analysis. Ectopically migrating cells from rosette were manually screened. Cell track speed, track length and displacement length for each cell were obtained using the Imaris program. The top 10 cells (ranked by absolute value of cell track speed, track length and displacement length, respectively) in each location were chosen and their values were normalized by subtracting the average of last 5 cells (ranked by absolute value of cell track speed, track length and displacement length, respectively) in corresponding location group to minimize the non-autonomous cell motion. A minimum of 10 different locations from three independent experiments were used to quantify cell migration for each genotype.

**QUANTIFICATION AND STATISTICAL ANALYSIS**

Statistical analyses were performed on Prism (GraphPad, version 9.2.0) using Student’s t-test, one-way ANOVA and two-way ANOVA based on data distribution. The details of the statistical test used for each figure were shown in the corresponding figure captions.

**DATA AND CODE AVAILABILITY**

The original data supporting the figures in the paper are available on request from the corresponding author.

Supplement tables:

1. Table S1_List of DEG_D10
2. Table S2_Figure 2E_GO BP_result_down
3. Table S3_Figure 2E_GO BP_result_up
4. Table S4_List of DEG in 14 cluster
5. Table S5_List of DEG_HOPX- oRG vs HOPX+ oRG
6. Table S6_Figure 4B_GO BP
7. Table S7_Figure S4A_ KEGG

Supplement video:

1. D10 WT hCO migration
2. D10 FEZ1-null hCO migration
